# New multiscale characterisation methodology for effective determination of isolation-structure-function relationship of extracellular vesicles

**DOI:** 10.1101/2021.02.09.430523

**Authors:** Thanh Huyen Phan, Shiva Kamini Divakarla, Jia Hao Yeo, Qingyu Lei, Priyanka Tharkar, Taisa Nogueira Pansani, Kathryn G. Leslie, Maggie Tong, Victoria A. Coleman, Åsa Jämting, Mar-Dean Du Plessis, Elizabeth J. New, Bill Kalionis, Philip Demokritou, Hyun-Kyung Woo, Yoon-Kyoung Cho, Wojciech Chrzanowski

## Abstract

Extracellular vesicles (EVs) have been lauded as next generation medicines, but very few EV-based therapeutics have progressed to clinical use. Limited clinical translation is largely due to technical barriers that hamper our ability to mass-produce EVs, *i.e.* to isolate, purify and characterise them effectively. Technical limitations in comprehensive characterisation of EVs leads to unpredicted biological effects of EVs. Here, using a range of optical and non-optical techniques, we showed that the differences in molecular composition of EVs isolated using two isolation methods correlated with the differences in their biological function. Our results demonstrated that the isolation method determines the composition of isolated EVs at single and sub-population levels. Besides the composition, we measured for the first time the dry mass and predicted sedimentation of EVs. These parameters were shown to correlate well with the biological and functional effects of EVs on single cell and cell cultures. We anticipate that our new multiscale characterisation approach, which goes beyond traditional experimental methodology, will support fundamental understanding of EVs as well as elucidate the functional effects of EVs in *in vitro* and *in vivo* studies. Our findings and methodology will be pivotal for developing optimal isolation methods and establishing EVs as mainstream therapeutics and diagnostics. This innovative approach is applicable to a wide range of sectors including biopharma and biotechnology as well as to regulatory agencies.

## Introduction

Current medicine has only taken us so far in reducing disease and the tissue damage that it causes. Extracellular vesicles (EVs) have been hailed as the next generation of medicines. EVs are membrane-surrounded nanoscale structures secreted ubiquitously by cells. They contain multiple substances that influence the function of surrounding cells [1] [2]. Since EV composition reflects the composition of the parent cell, EVs are ideal candidates for use in disease diagnosis [3]. EVs are already considered as diagnostic biomarkers for cancer, cardiovascular, neurodegeneration, and kidney diseases [3–5]. It is also well-established that EVs transfer genetic information between cells and influence the behaviour of surrounding cells [6]. Therefore, they can be used for therapeutic applications such as tissue repair. EVs derived from stem cells are recognised as ‘second generation’ stem cell therapies [3, 7] – made by cells for cells. Compared to stem cells, EVs have key advantages including low immunogenicity, no ability to self-replicate (no risk of cancer), high resistance to hostile environments and improved bioactivity and stability upon storage [8]. However, despite all these potential advantages, very few EV applications have progressed to clinical use [9].

The limited clinical translation of EVs is largely due to technical barriers that hamper the mass-production of EVs, *i.e.* the ability to isolate, purify and characterise them effectively at single vesicle (nano), sub-population and population levels [10]. Since EV sizes range from 50 to 150 nm and they are secreted into rich multicomponent media or body fluids, isolation and characterisation are not trivial and remain as key challenges in the field [10]. The molecular corona that forms on EVs adds to the complexity of these challenges [11]. Since the corona changes the physicochemical characteristics of EVs and their affinity to the substrates used in some isolation methods, it is difficult to isolate and characterise EVs effectively. Moreover, EV isolates often contain lipoproteins, protein aggregates and non-vesicle macromolecules [12]. We also know that cell-free DNA can adsorb to lipid nanoparticles [13], which is likely to occur for circulating EVs too, but surprisingly this phenomenon is largely overlooked in the field. These ‘contaminants’ influence the biological function of EVs and are not trivial to detect due to the sensitivity of experimental methods. This also means that it is difficult to decouple them from the isolated EV populations [14]. However, these contaminants could potentially work synergistically with EVs to achieve specific therapeutic function in the body [15]. Therefore, for practical utilisation of isolation protocols it is necessary first to perform comprehensive physicochemical and molecular characterisation of EV isolates at sub/population and single vesicle levels and to measure functional responses to EVs in adequate cell/animal-based models.

The most commonly used approach to isolate EVs is ultracentrifugation, which involves multistep differential centrifugations to pellet vesicles [16, 17]. However, ultracentrifugation is labour-intensive, requires large sample volumes and produces a relatively low yield of enriched EVs [18]. Numerous alternative isolation methods have been developed including density gradients (DG) and size exclusion chromatography (SEC). Although DG and SEC usually result in high purity EV, these protocols are time-consuming, characterised by poor yields and suitable for small input volumes only (less than 5 mL) [19, 20]. Immunoaffinity-based approaches can also be used for EV isolation [21]. In these approaches, EVs are ‘collected’ by beads functionalised with EV-specific antibodies, thus isolation process is solely related to affinity of EVs to selected antibodies. The collection of EVs based on the assumption that specific markers are present on the surface of EVs. This means that these approaches are highly selective and likely to isolate only some fractions of EVs populations. More importantly, these methods fail to account for corona which cloaks EVs and is likely to cover some of the surface markers making these approaches even more selective [11]. Furthermore, at this stage only small quantities of biological samples can be processed in immunoaffinity-based isolation [22]. In contrast, tangential flow filtration (TFF) is capable of processing scalable volumes of biological fluids and producing high EV yield [23]. TFF is technically simple to operate and requires low-cost instrumentation. These features make TFF well-suited to isolate EVs at large scale. However, to enable broader applications of TFF, its advantages in comparison with ultracentrifugation (and other methods) must be determined.

Previous studies investigated the differences in physical properties between EVs isolated using TFF and ultracentrifugation [23, 24]. However, the results are limited to selected physical characteristics of EVs (e.g., yield, size distribution, morphology, surface markers), and do not show their correlation with biological effects. To the best of our knowledge, there is no available studies investigating how the physicochemical properties of EVs contribute to the EV functionality. Here, using a combination of high-resolution optical and non-optical techniques, as well as functional assays, we interrogated the differences between EVs isolated using ultracentrifugation and TFF at single vesicle, sub-population and population levels. The significance of this work is in the demonstration of how the isolation methods influence the physicochemical/molecular composition of EVs and functional cell responses to EVs. For the first time we measured the dry mass of large EVs or EV agglomerates (>100 nm), predicted the sedimentation of EVs and developed a new probe to assess the functionality of EVs. The key strength of this study lies in the comprehensive characterisation of EVs at single vesicle, EV sub-population and population levels, as well as in the analysis of cellular responses *(i.e.* EV uptake) at single cell level. To achieve desired statistical and scientific validation of our approach we used two cell types which produce different quantities of EVs and that these EVs characterise with different molecular cargo; specifically we used chorionic and decidual mesenchymal stem cells (CMSC29 and DMSC23 respectively) [25, 26].

Our study provides evidence that EVs’ physicochemical characteristics (at single vesicle, subpopulation and population level) as well as their biological function, depends on the isolation method. The differences in physicochemical properties of EVs were shown to correlate well with the functional effects of EVs. We concluded that isolation method determines the composition of EV isolates. These findings are of critical significance in the field because they suggest that isolation method is pivotal in establishing downstream applications of EV as diagnostic biomarkers, therapeutics and in fundamental biology. Notably, this work highlights the importance of nanoscale and single particle characterisation methods in EV research and the need for the simultaneous and integrated use of physicochemical and functional assays.

## Results

To measure EVs size and concentration we used Particle Tracking Analysis (PTA), Dynamic Light Scattering (DLS) (size only), Nano-flow Cytometry (nFCM), Tunable Resistive Pulse Sensing (TRPS), and Asymmetric Flow-Field fractionation (AF4) (size only). Nanoscale infrared spectroscopy (AFM-IR) and nFCM were used to determine EV composition at the single EV and EV sub-population levels. For the first time, we used Resonant Mass Measurement (RMM) for the characterisation of dry mass and buoyant mass of large EVs (>100 nm) and distorted grid (DG) for the sedimentation prediction of EVs. Sedimentation of nanoparticles reveals the actual concentration of nanoparticles on the cell surface, which correlates with cellular uptake [23]. Therefore, sedimentation is the key factor in the interpretation of downstream biological effects of EVs on the cellular response. The actual functional effects of EV isolates on cells was determined using newly developed nitric oxide fluorescent probe to measure intracellular stress in an *in vitro* model of acute lung injury.

### Comparison of EV size distribution and concentration

Chorionic and decidual MSCs cell lines (CMSC29 and DMSC23) were used as cell sources to isolate EVs. EVs isolated from CMSC29 and EVs isolated from DMSC23 cells were referred to as CEVs and DEVs, respectively.

The size distribution of CEVs and DEVs was assessed using four independent techniques; PTA, DLS, nFCM and TRPS.

### PTA

PTA analyses showed that both CEVs and DEVs isolated using TFF and ultracentrifugation had a similar particle size distribution ranging from 100 to 300 nm **(Fig. 1 A, B)**. However, there was a small peak at around 50 nm in the size distribution spectrum of DEVs isolated using TFF, which was not present in EVs isolated using ultracentrifugation **(Fig. 1 B)**.

**Fig. 1.**
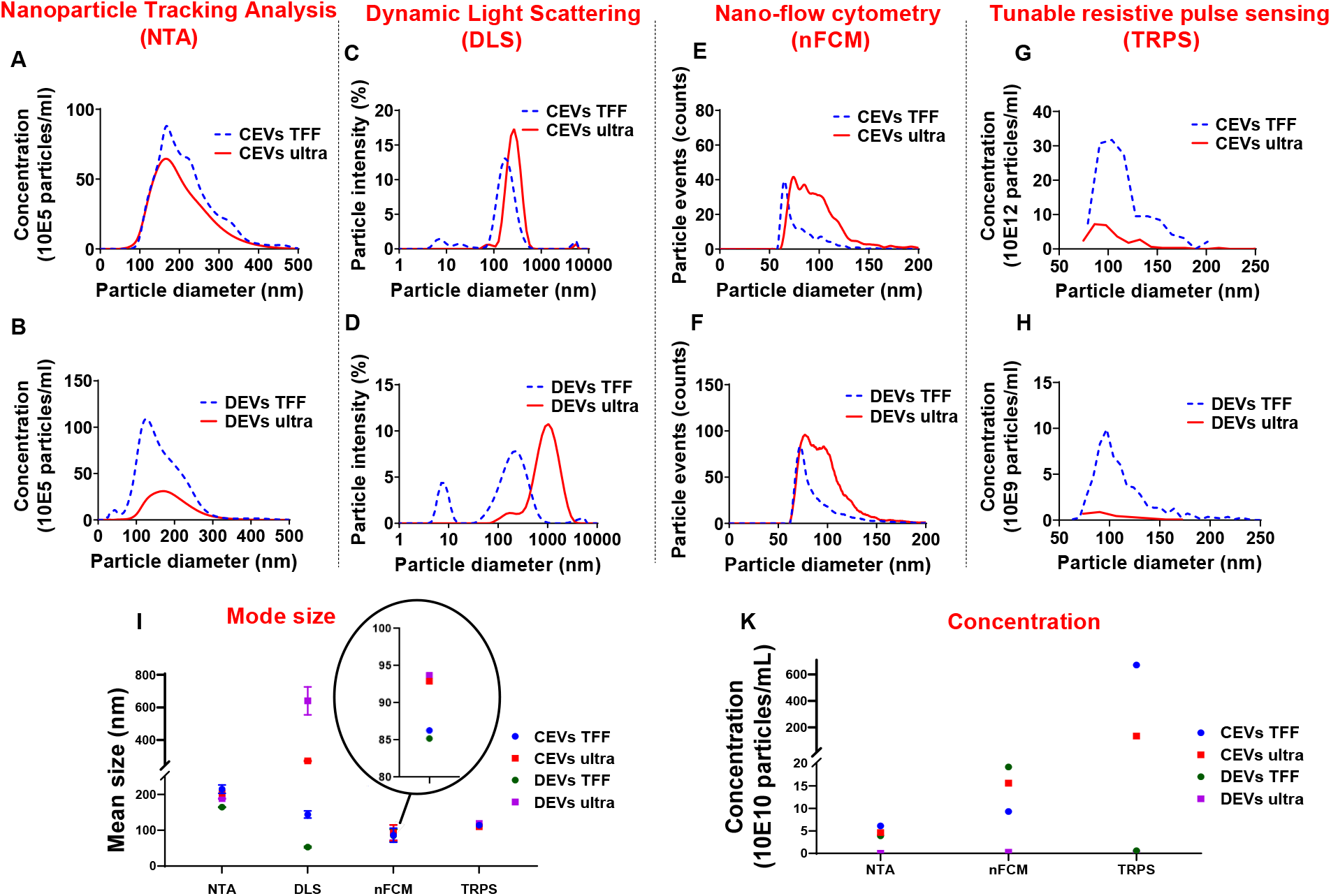
Comparison of the size distribution for different isolated EVs. Representative histograms revealing size distribution of CEVs and DEVs isolated using TFF and ultracentrifugation by 4 different methods. The size distributions of these EVs were assessed by (**A**) and (**B**) Particle Tracking Analysis (PTA), (**C**) and (**D**) Dynamic Light Scattering (DLS), (**E**) and (**F**) Nano-flow Cytometry (nFCM) and (**G**) and (**H**) Tunable Resistive Pulse Sensing (TRPS) respectively. The mode size **(I)** and concentration **(K)** of all EV samples were presented.

### DLS

Size analyses using DLS showed that the intensity-based size distribution of CEVs and DEVs isolated using TFF and ultracentrifugation were different **(Fig. 1 C, D)**. CEVs isolated using TFF had one high-intensity peak at around 200 nm with one low-intensity peak at approximately 8 nm. In contrast, CEVs isolated using ultracentrifugation had one dominant peak at around 300 nm **(Fig. 1 C)**. The intensity-based size distribution for DEVs isolated using TFF had two peaks at 8 nm and 200 nm, while DEVs isolated using ultracentrifugation had one broad peak at around 1000 nm with one small shoulder at around 200 nm **(Fig. 1 D)**.

The presence of small particles (around 8 nm) in CEVs isolated using TFF was verified using preliminary asymmetric flow-field fractionation (AF4) measurement **(Fig. S2)**. Since the elution peak of CEVs isolated using TFF at 18 min coincides with pure BSA, the small-size particles in CEVs isolated using TFF were identified as BSA contaminants.

Overall, the size of both CEVs and DEVs isolated using ultracentrifugation as assessed by DLS were larger than the EVs isolated using TFF. This is likely to be related to the presence of a few larger entities (potentially agglomerates during the ultracentrifugation process). Since DLS is extremely sensitive to the presence of large entities, even with a low amount of these would account for the shape of the intensity-weighted data [27].

### nFCM

nFCM measurements showed a broad size distribution ranging from 50 to 200 nm for both CEVs and DEVs isolated using TFF and ultracentrifugation **(Fig. 1 E, F)**. While the size of most of EVs isolated using TFF (for both CEVs and DEVs) was around 60 nm, the size of EVs isolated using ultracentrifugation was evenly distributed from 50 to 200 nm.

### TRPS

TRPS measurements showed that CEVs isolated using TFF and ultracentrifugation had a similar size distribution spectrum **(Fig. 1 G)**. The size of DEVs isolated using TFF showed a broad distribution ranging from 70 to 200 nm while the size distribution obtained for DEVs isolated using ultracentrifugation could not be used for comparison due to a much lower sample concentration **(Fig. 1 H)**.

The mode sizes of both CEVs and DEVs were consistent for each individual method (with differences between methods) regardless of the isolation method. PTA, nFCM and TRPS showed the mode sizes of CEVs and DEVs were around 150, 90 and 110 nm respectively (**Fig. 1 I**). However, DLS showed differences in mode sizes of CEVs and DEVs depending on the isolation method. Ultracentrifugation yielded larger CEVs with mode size was around 270 nm, while CEVs isolated with TFF was approximately 197 nm. Similarly, the mode size of DEVs isolated using ultracentrifugation was approximately 1120 nm, and TFF isolated DEVs was approximately 240 nm. It is important to notice that the scattering intensity depends on the 6^th^ power of the size of macromolecules, therefore, large agglomerates-even a very small amount will overwhelm the intensity from small size particles in DLS [28]. Thus, the observation of small size particles in EVs isolated using TFF in DLS indicated the substantial amount of small size particles and the absence of agglomerates in TFF isolated EVs. To further explore the possibility of the presence of small size particles in the samples, AF4 was used **(Fig. S2)** to fractionate and measure the hydrodynamic diameter of the entities as they eluted.

The concentration of EVs was determined using three techniques: PTA, nFCM and TRPS **(Fig. 1 K)**. The concentration of CEVs isolated using TFF and ultracentrifugation determined by PTA and nFCM was approximately 10^11^ EVs/mL. In contrast, TRPS measured approximately 10^12^ CEVs/mL isolated using TFF, one order of magnitude larger than PTA and nFCM. The concentration of DEVs isolated using TFF measured using PTA and TRPS was ~10^9^ EVs/mL, while the measurement made using nFCM was 10^11^ EVs/mL, two orders of magnitude higher. The concentration of DEVs isolated using ultracentrifugation was consistent for all measurement methods, 10^9^ EVs/mL.

Although multiple studies have implemented PTA and DLS as standard techniques to measure EV sizes, both methods have some limitations. PTA is based on light scattering and Brownian motion, which cannot differentiate between large protein aggregates and single large particle [29]. For DLS, the size distribution depends on the intensity of light scattering, therefore the intensity of the scattered light from small particles is normally surpassed by large particles [30]. In addition, many EVs have protein corona on their surfaces, which may impact on the accuracy of the results as EVs with and without corona (or unspecific corona) have different hydrodynamic diameter [31]. Therefore, DLS provides only information about relative and not accurate size. The nFCM has higher sensitivity compared to PTA due to its lower detection threshold. Indeed, there are changes to the angles of the light scatter collected by nFCM, which allow the detection down to 40 nm [32]. However, nFCM was calibrated to detect particles below 200 nm, therefore the agglomerates could not be detected using this technique. TRPS detects vesicles within the range of the nanopore selected [33]. As nanopore NP100 (50-330 nm size range) was used in this study, TRPS was less sensitive for detection of EV particles below 50 nm. Size and concentration measurement results showed that each characterisation technique has its own limitations due to principles of the measurement, the way samples are prepared as well as calibration standards (for nFCM and TRPS). Consequently, as evidenced here, it led to discrepancies between the data generated from each of the techniques. These findings emphasize the need to use multiple complementary techniques. The interpretation of the data must take into account the principle behind the measurement technique, as this can affect the results.

In summary, both PTA and nFCM measurements showed a consistent average size for both DEVs and CEVs regardless of isolation method, however the concentration of EVs was around two orders of magnitude lower for DEVs isolated using ultracentrifugation. Overall, results suggested that PTA was less sensitive than nFCM for the detection of EVs smaller than 100 nm.

### Mass analysis

Besides the size distribution, the buoyant mass and dry mass of different isolated EVs were quantified. Previously, buoyant mass and dry mass were used to determine the exact amount of nanoparticles interacting with cells and tissues in toxicity studies [34]. Here, we used resonant mass measurement (RMM) for the first time to characterise EVs for buoyant mass and dry mass. The advantage of this approach is that both mass parameters can be measured for individual vesicles which in turn reveals the total mass, including molecular cargo, of each vesicle [35]. Precise knowledge of the amount of EV cargo is pivotal to define the biological function of EVs.

The dry mass of EVs isolated using ultracentrifugation and TFF was measured and presented in **Fig. 2 A, B.** The size distribution of each EV type was calculated from the measured buoyant mass, with a particle density of 1.4 g/cm^3^ **(Fig. S3)**. It is important to notice that the limit of detection (LOD) of RMM for dry mass is 10^-15^ g and for size distribution is 100 nm. Therefore, only sub-population of EVs with the high mass was detected and measured. When plotted as a function of dry mass, both detectable CEVs and DEVs isolated using ultracentrifugation showed higher dry mass than EVs isolated using TFF **(Fig. 2)**. The dry mass of measurable CEVs isolated using ultracentrifugation (4.67 × 10^-15^ g) was higher than CEVs isolated using TFF (3.17 × 10^-15^ g) **(Fig. 2 A)**. Similarly, measurable DEVs isolated using ultracentrifugation (7.43 × 10^-15^ g) showed higher dry mass than DEVs isolated using TFF (3.9 × 10^-15^ g) **(Fig. 2 B)**. Interestingly, a sub-population of DEVs isolated using TFF which has the positive buoyancy was detected and its size was estimated using a density of 0.01 g/cm^3^ **(Fig. S3)**. This positively buoyant sub-population of DEVs isolated using TFF could be bubbles or empty vesicles. The higher mass for detectable EVs (above 100 nm) (CEVs and DEVs) isolated using ultracentrifugation suggested the presence of more agglomerates in ultracentrifugation isolated EVs.

**Fig. 2.**
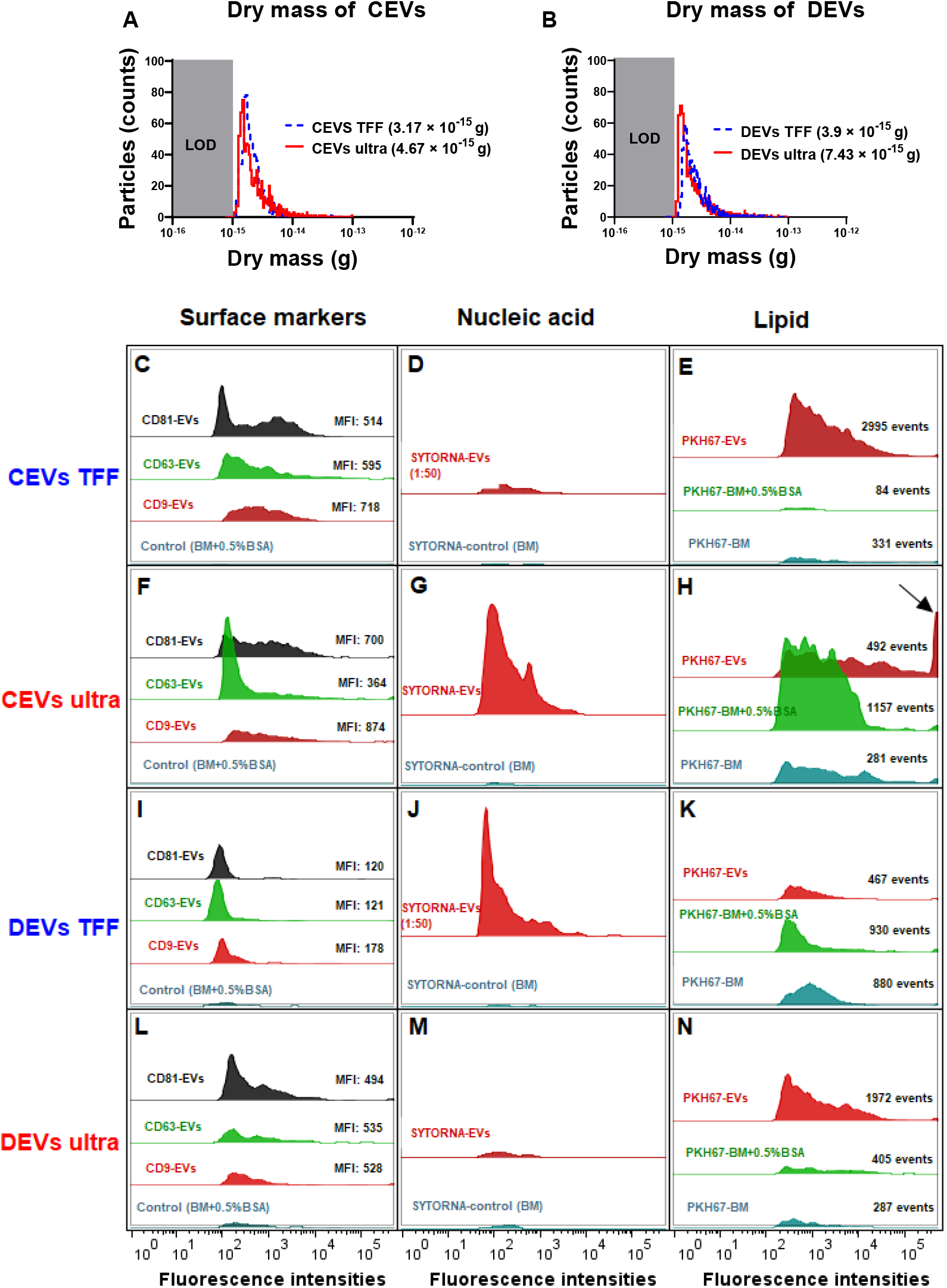
The dry mass of different isolated EVs and EV molecular composition was analysed using nFCM. The dry mass of TFF and ultracentrifugation isolated **(A)** CEVs and **(B)** DEVs. Error bar in mode size of EVs denotes the standard deviation (SD). LOD: limit of detection. Representative histograms showing the presence of three surface markers CD9, CD63 and CD81 **(C, F, I, L)**, nucleic acid **(D, G, J, M)**, lipid content **(E, H, K, N)** in CEVs and DEVs isolated using TFF and ultracentrifugation. The arrow indicates the saturated fluorescence peak. MFI: Mean fluorescence intensity.

Cumulatively, size and mass measurements suggested that these EVs contained some amount of agglomerates. The agglomeration of EVs during ultracentrifugation was reported in previous studies [36–38].

### EV molecular composition assessed using nFCM and nanoscale infrared spectroscopy (AFM-IR)

#### EV surface markers

The differences in dry mass of EVs isolated using ultracentrifugation and TFF suggested that there were differences in molecular composition of EVs. Therefore, we assessed the presence of three common EV surface markers (i.e. CD9, CD63 and CD81) on different isolated EVs by using nFCM. CD9, CD63, CD81-positive EVs were defined as the events that were above the threshold level and detected by fluorescence triggering **(Fig. S4)**. A control using basal medium with 0.5% (w/v) BSA **(Fig. S4)** showed negligible fluorescence in fluorescence triggering, which demonstrated that the free dye was successfully removed by our washing procedure. The histogram analyses showed that both CEVs and DEVs regardless of isolation method were positive for CD9, CD63 and CD81 **(Fig. 2 C, F, I, L)**. The overall mean fluorescence intensity (MFI) of CD9, CD63 and CD81 was higher for CEVs isolated using the same method. Profile analysis of individual surface markers showed that their expression was not uniform across different sub-populations of EVs. CD9 was detected as strongly positive in CEVs isolated using TFF (MFI: 718), followed by CD63 (MFI: 595) and CD81 (MFI: 514) **(Fig. 2 C)**. Whereas, the expression of CD9 (MFI: 874) and CD81 (MFI: 700) was higher than CD63 (MFI: 364) in CEVs isolated using ultracentrifugation **(Fig. 2 F)**.

The expression of the markers was more uniform for DEVs than CEVs when using the same isolation method. All three markers of DEVs isolated using TFF were uniformly detected at a low MFI level (under 200) **(Fig. 2 I)**. Specifically, the MFI value for CD9 (178) was higher than CD63 and CD81 (121 and 120 respectively). Since DEVs isolated using TFF had substantial amount of small size particles (~8 and 200 nm) assessed using DLS, the expressions of the markers were lowest. Meanwhile, DEVs isolated using ultracentrifugation showed higher expression of the markers than TFF. The MFI values for CD9, CD63 and CD81 were 494, 535 and 528 respectively in DEVs isolated using ultracentrifugation **(Fig. 2 L)**.

The profile of the surface markers for EVs isolated from each of the cell type was different depending on the isolation method. This result suggested that each of the isolation method provided different EV populations. However, regardless of the isolation method EVs isolated from DMSC23 consistently showed more uniform expression of all three markers, which implied that exosome-specific markers were more homogenously expressed on DEVs.

#### Nucleic acid profiling

We next quantify the total nucleic acid content inside EVs by staining EVs with SYTO RNASelect green fluorescent cell stain. The percentage of nucleic acid was calculated by dividing the concentration of SYTO RNASelect green positive events by the total concentration of EVs **(Table 1)**. EVs isolated using TFF were diluted 50 times before staining to achieve the same threshold level as EVs isolated using ultracentrifugation. Since TFF isolated EVs were diluted, a smaller fluorescence intensity peak was shown in CEVs isolated using TFF in comparison to CEVs isolated using ultracentrifugation **(Fig. 2 D, G)**. However, the analyses after the calculation showed that CEVs isolated using TFF contained 22 times more of the nucleic acid (2.46%) than CEVs isolated using ultracentrifugation (0.11%) **(Table 1)**. Similarly, the nucleic acid content of DEVs isolated using TFF (3.31%) was 7 times higher than DEVs isolated using ultracentrifugation (0.46%) **(Table 1, Fig. 2 J, M)**. Overall, we concluded that the total amount of nucleic acid in DEVs was higher than CEVs when using the same isolation method.

**Table 1.** The percentage of nucleic acid content in CEVs and DEVs isolated using TFF and ultracentrifugation assessed using nFCM.

| EV type | Percentage of nucleic acid (%) |
| --- | --- |
| CEVs TFF | 2.46 |
| CEVs ultra | 0.11 |
| DEVs TFF | 3.31 |
| DEVs ultra | 0.46 |

#### Lipid membrane profiling

We next assessed the lipid content in each EV type by staining EVs with green fluorescence lipophilic dye, PKH67. PKH67, which labels the cell membrane by inserting its aliphatic chains into the lipid membrane, has been used extensively to label the lipid membrane of EVs [39]. The lipid compositions of DEVs and CEVs were measured and compared with two controls: basal medium only (BM) and basal medium with 0.5% (w/v) BSA (BM+0.5% BSA) **(Fig. 2)**. We used two controls as PKH67 may label other components in medium (non-specific binding) [40, 41]. Hence, two control groups were essential to eliminate false positives due to fluorescence from non-EV components.

The total fluorescence events of CEVs isolated using TFF (2995 events) was 9 and 35-fold higher than BM (84 events) and BM+0.5% BSA (331 events) **(Fig. 2 E)**, which showed substantial PKH67-positive EVs. On the other hand, CEVs isolated using ultracentrifugation (492 events) had 221 more fluorescence events than BM (281 events), but they had 665 less events than BM+0.5% BSA (1157 events) **(Fig. 2 H)**. The fluorescence positive events in controls indicated that there were some ‘contaminants’ in controls, which bound to PKH67 and caused the fluorescence. The saturated fluorescence intensity peak in CEVs isolated using ultracentrifugation, which was not detected in controls, suggested the presence of larger size vesicles or agglomerates **(Fig. 2 H; arrow)**.

The total fluorescence events of DEVs isolated using TFF (467 events) were approximately two-fold lower than BM (880 events) and BM+0.5% BSA (930 events) **(Fig. 2 K)**. DEVs isolated using ultracentrifugation (1972 events) had 1685 and 1567 more events than BM (287 events) and BM+0.5% BSA (405 events) respectively **(Fig. 2 N)**. This result indicated that DEVs isolated using ultracentrifugation contained high concentration of PKH67-positive EVs. Since DEVs isolated using ultracentrifugation had the lowest concentration of EVs among all EV groups, this result could be a false positive result. One possible explanation could be that during ultracentrifugation lipid-protein aggregates and other media components are isolated beside EVs. These undesired components can potentially bind PKH67, which causes a misleading result.

As observed in **Fig. 2 H** and **Fig. 2 N**, the number of events shown in the controls (BM and BM+0.5% BSA) emphasized the non-selective binding of PKH67 to other components. These results are consistent with the findings that the PKH67 lipophilic dyes is not specific to EVs [40]. Therefore, this study was inconclusive regarding to the amount of lipid of EV isolates.

#### Molecular composition of individual EVs using nanoscale infrared spectroscopy (AFM-IR)

While nFCM proved to be a very effective method to determine size and composition of EVs at bulk population level, this method, similar to other size measurements methods, cannot effectively distinguish between a single large EV and an agglomerate of several small EVs [36]. Therefore, to characterise EVs at single vesicle level we used nanoscale infrared spectroscopy (AFM-IR) **(Fig. 3)** [42, 43]. Topographical AFM images showed that both CEVs and DEVs were spherical and their size ranged between 20-300 nm regardless of isolation method **(Fig. 3 A)**. Molecular analysis at the single vesicle level employed state-of-the-art nanoscale infrared spectroscopy (AFM-IR), which showed that individual EVs contained proteins, nucleic acid and lipids **(Fig. S5, Fig. 3 B, C)**. However, the compositions of both types of EVs were different depending on the isolation method.

**Fig. 3.**
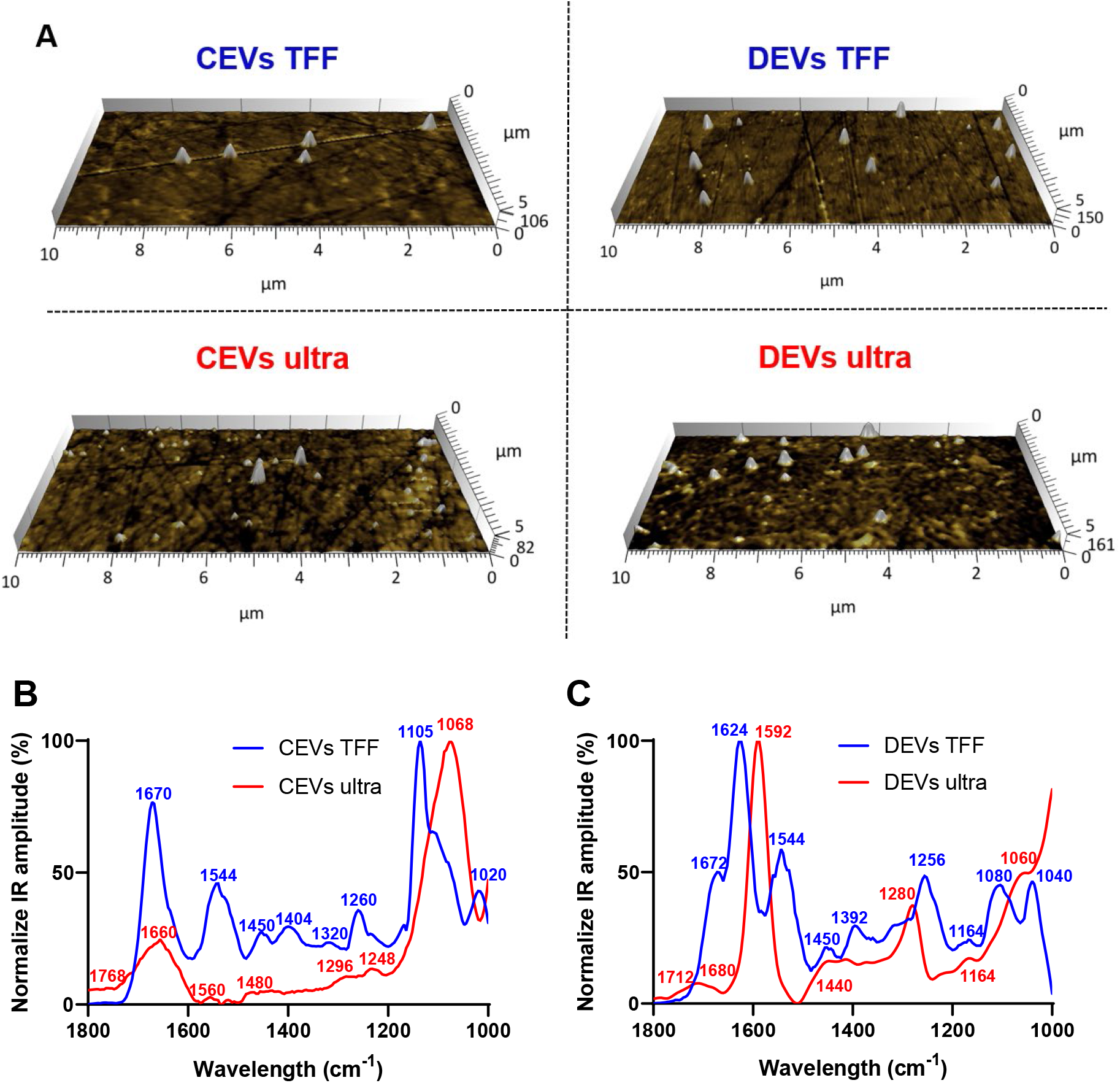
Average infrared absorption spectra of EV molecular composition corresponding topographical AFM images for EVs. **(A)** AFM images (10 × 5 μm) of individual EVs (indicated as white protrusions) deposited on prism. Normalization of the average AFM-IR spectra of **(B)** CEVs isolated using TFF and ultracentrifugation; **(C)** DEVs isolated using TFF and ultracentrifugation. At least 15 individual EVs were randomly selected and analysed using Analysis Studio™ software.

CEVs isolated using TFF had dominant peaks for all three main components of EVs; protein (1670, 1544, 1260 cm^-1^), nucleic acid (1450, 1404, 1320 cm^-1^) and lipid (1105, 1020 cm^-1^), which confirmed the presence of all three component in individual EVs [44]. The intensity ratio of protein amide I (1670 cm^-1^) and amide II (1544 cm^-1^) was 2:1, which indicated ordinary protein conformation in CEVs isolated using TFF. Peaks at 1450 and 1404 cm^-1^ in CEVs isolated using TFF spectra were attributed to phosphatidylcholine head group and thymine of RNAs [43]. In contrast, CEVs isolated using ultracentrifugation had a broad band with low intensity in amide I peak at 1660 cm^-1^. They also showed a dominant peak at 1068 cm^-1^ (lipid) and five minor peaks at 1768, 1560, 1480, 1296 and 1248 cm^-1^. Since the intensity of amide II peak (1560 cm^-1^) was low in CEVs isolated using ultracentrifugation, it suggested that protein conformation for these EVs was altered. Moreover, the absence of peaks in the 1450-1350 cm^-1^ region in CEVs isolated using ultracentrifugation suggested smaller amounts of nucleic acid (i.e. RNAs), which was consistent with nFCM result **(Table 1)**. The dominant peak at ~1068 cm^-1^ in CEVs isolated using ultracentrifugation spectra was associated with the ester C–O–C symmetric stretching vibration [45]. This peak was shifted to ~1105 cm^-1^ in CEVs isolated using TFF spectra. The peak at 1768 cm^-1^ in CEVs isolated using ultracentrifugation spectra could be related to ester groups, primarily from lipid and fatty acids [42].

DEVs isolated using TFF characterized showed dominant peaks at 1624, 1544, 1256 cm^-1^ and a shoulder at 1672 cm^-1^ (protein), 1450, 1392 cm^-1^ (nucleic acid) and 1080, 1040 cm^-1^ (lipid). The intensity ratio of amide I (1624 cm^-1^) and amide II (1544 cm^-1^) peaks was ~2:1, which suggested unaltered protein conformation in DEVs isolated using TFF. Peaks at 1450 and 1392 cm^-1^ in DEVs isolated using TFF spectra confirmed the presence of RNAs [43]. In contrast, DEVs isolated using ultracentrifugation had dominating peaks at 1592 cm^-1^ (protein amide II), 1280 cm^-1^ (nucleic acid, protein) and broad bands at 1680, 1440 and 1164 cm^-1^. The low intensity of the amide I (1680 cm^-1^) peak in DEVs isolated using ultracentrifugation suggested some alterations in protein structures of these EVs. The broad band with low-intensity at 1440 cm^-1^ in the spectra of DEVs isolated using ultracentrifugation suggested a reduced amount of RNAs, which was consistent with the nFCM result **(Table 1)**. In addition, the bands in the 1080-1040 cm^-1^ region were associated with vibration from phosphate stretch of RNA/DNA or lipid [46]. While DEVs isolated using TFF had two peaks at 1080 and 1040 cm^-1^, DEVs isolated using ultracentrifugation only had one peak at 1060 cm^-1^ suggesting changes in RNA/DNA structures and lipid. The peak at 1712 cm^-1^ in the spectra of DEVs isolated using ultracentrifugation could be related to ester groups, primarily from lipid and fatty acids.

Overall, the structures of all three-key molecular/structural components of EVs, proteins, lipids and nucleic acid were maintained in both CEVs and DEVs isolated using TFF. In contrast, for EVs isolated using ultracentrifugation the intensity, position and the width of the peaks changed, indicating changes to the molecular structure of these components. These changes were observed for both CEVs and DEVs.

Since both nucleic acid and lipid peaks are centred around the same frequency it was necessary to validate the results using an alternative technique, e.g. nFCM. We showed that the amount of nucleic acid in EVs isolated using ultracentrifugation was smaller than TFF regardless of the origin of EVs, which was consistent with the nFCM result. Therefore, combining both nFCM and nanoscale infrared spectroscopy are necessary to gain precise understanding of the molecular/structural composition of individual EVs.

#### Predicted sedimentation of isolated EVs using distorted grid (DG) model

The differences in physicochemical properties of EV types suggested that their colloidal stability, interactions with the cell membrane, internalisation and downstream biological effects could also be different [47]. Whilst largely ignored, the colloidal stability is a critical parameter that shows ability of EVs to move towards, and reach cells (‘sediment’), hence defining biological effectiveness of EVs. The transport modelling for EVs was completed for the first time based on their size, size distribution, effective density to calculate the particles sedimentation and diffusion in cell culture media.

The predicted sedimentation of different EV isolates was assessed using the DG model. All the values used for the modelling are shown in **Table S2**. The results indicated the differences in sedimentation between EVs depending on the isolation method **(Fig. 4)**. CEVs isolated using TFF showed the deposited fraction was predicted to reach 0.00354 within 1 h, then slowly reach the mean fraction deposited (0.003587) after ~10 h **(Fig. 4 B)**. Meanwhile, CEVs isolated using ultracentrifugation was predicted to reach the mean fraction deposited (0.2169) after ~7.5 h **(Fig. 4 D)**. For DEVs, the mean fraction deposit was predicted to be reached after ~8 h for EVs isolated using TFF (0.003588) **(Fig. 4 C)** and ~10 h for EVs isolated using ultracentrifugation (0.3198) **(Fig. 4 E)**. Overall, the mean of fraction deposit of EVs isolated using ultracentrifugation was 65 and 100 times higher than EVs isolated using TFF for CEVs and DEVs respectively. A previous study reported that less agglomeration resulted in a smaller deposited fraction [48], which confirmed that EVs isolated using TFF do not contain agglomerates. In contrast, EVs isolated using ultracentrifugation may contain the agglomerates and these agglomerates attributed to the higher sedimentation. This result was consistent with our size distribution (DLS), mass (RMM) and visualization EVs (holotomography) findings.

**Fig. 4.**
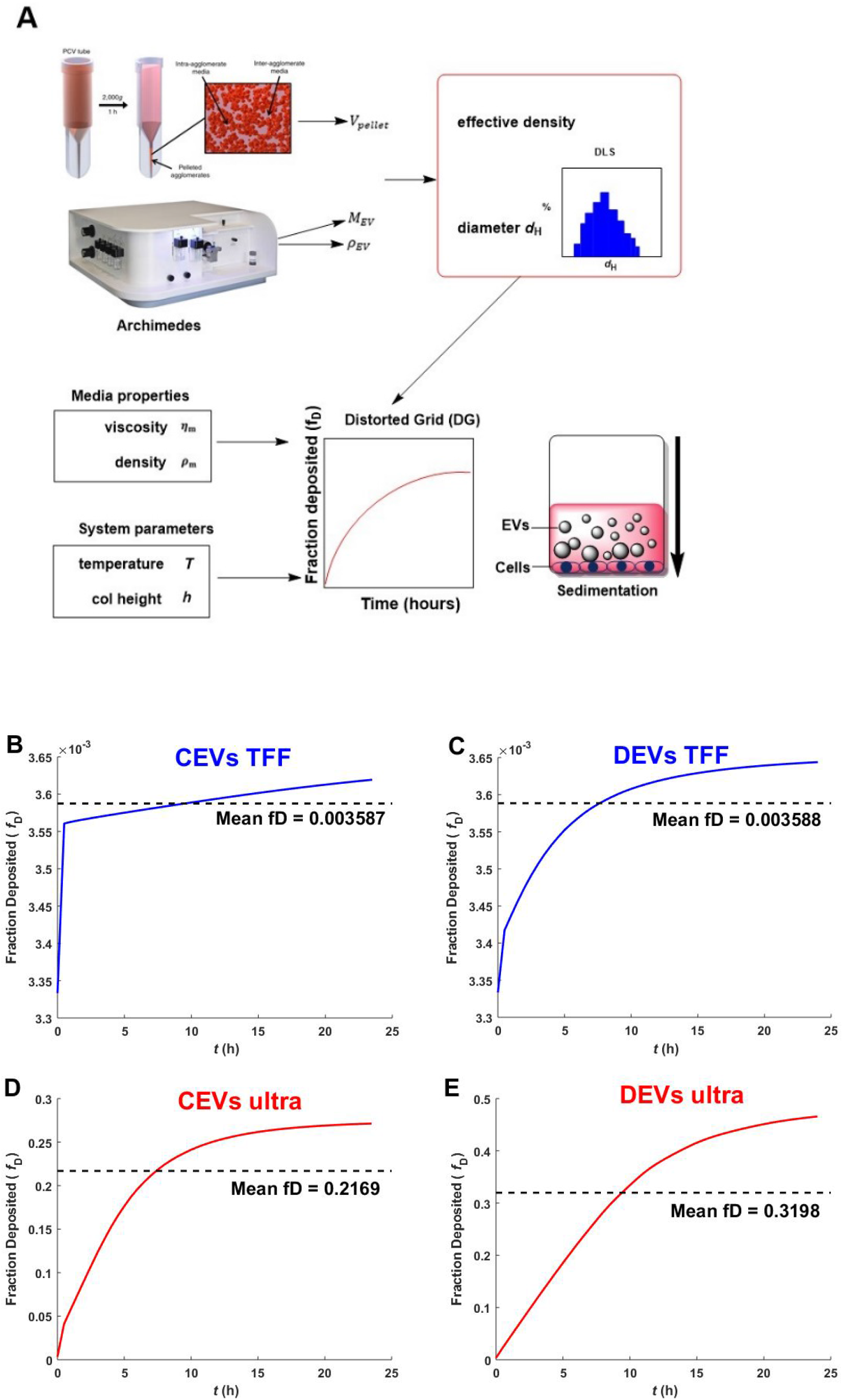
Schematic overview of nanodosimetry approach and the predicted fraction deposited of EVs. The schematic overview of DG modelling adapted from Deloid *et al.* [49] **(A)**. The predicted fraction deposited of CEVs isolated using **(B)** TFF and **(D)** ultracentrifugation; DEVs isolated using **(C)** TFF and **(E)** ultracentrifugation into each well in 96 well plate was generated by Matlab. Mean fD is the mean fraction deposited.

#### EV internalisation by living BEAS-2B cells using holotomography

To confirm the predicted sedimentation of EVs, the uptake of EVs isolated using TFF and ultracentrifugation by cells was visualised using correlative holotomography and fluorescence microscopy **(Fig. 5)**. This method, unlike confocal microscopy, does not require any fluorescent labelling for cells, which imparted no stress on cells. Additionally, holotomography microscopy allows us to visualize the differences in refractive index, thus we can observe sub-cellular organelles of the cells without labelling [50].

**Fig. 5.**
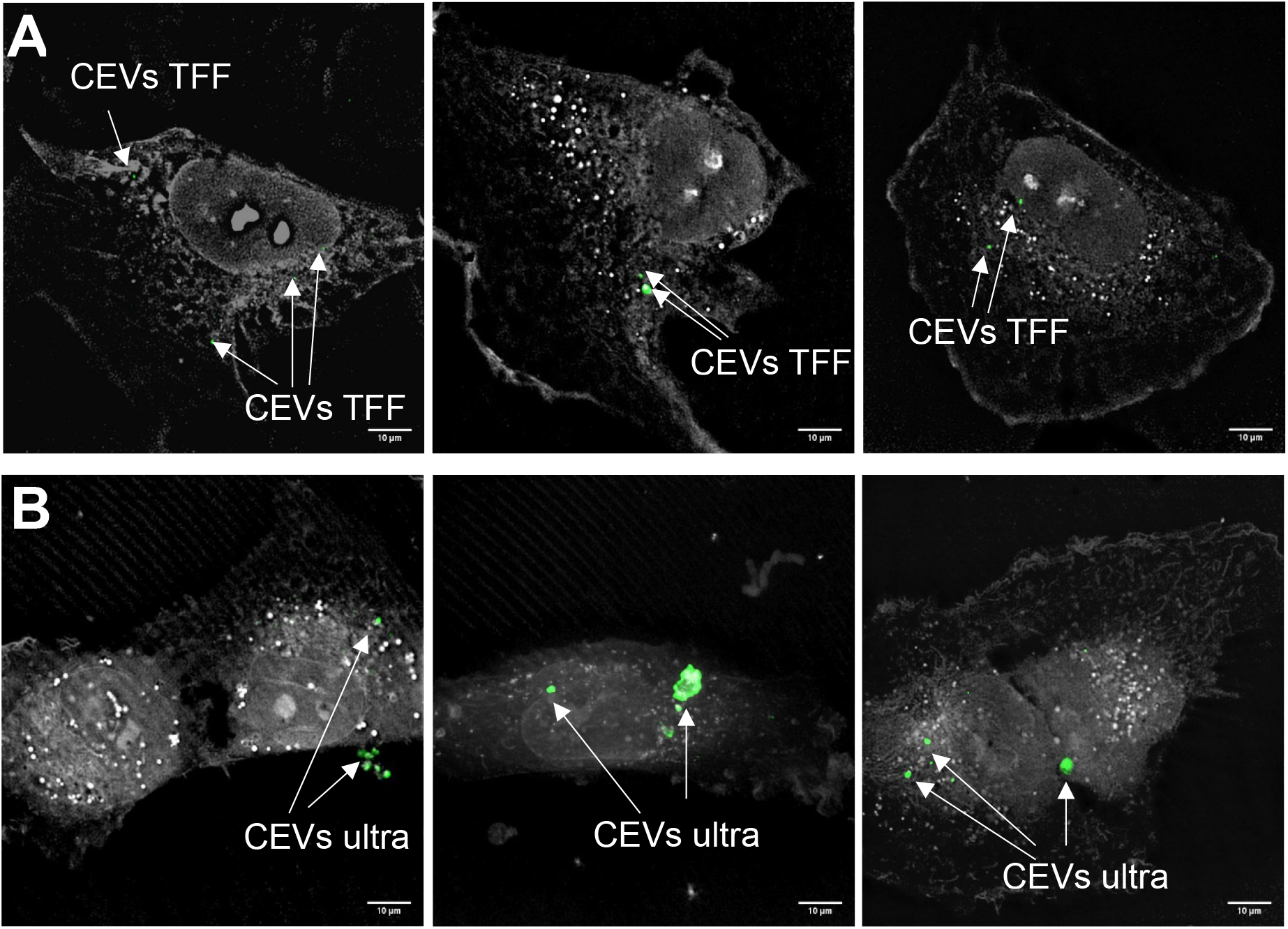
The localisation of EVs with BEAS-2B cells. Three representative images of BEAS-2B cells and stained CEVs (green colour) isolated using **(A)** TFF and **(B)** ultracentrifugation. Scale bar represents 10 μm. Bright white dots inside BEAS-2B cells indicate the lipid droplets.

DEVs isolated using TFF had substantial amount of small size particles **(Fig. 1)**, which caused the difficulties in visualising EVs. Therefore, to demonstrate the differences between TFF and ultracentrifugation isolated EVs, the EVs localisation study using a holotomography microscope was conducted only for CEVs. BEAS-2B cells were dosed with PKH67-labelled CEVs and imaged 3 h later. We observed fluorescence inside the cells and not on the plasma membrane, which suggested that EVs were internalised by the cells. The results showed that the punctate CEVs isolated using TFF and ultracentrifugation were qualitatively homogenously distributed in the cytoplasm **(Fig. 5 A)** and of uniform size. The size of CEVs isolated using ultracentrifugation were heterogenous, ranging from 100 nm to 500 nm **(Fig. 5 B)**. These observations confirmed agglomeration had occurred during ultracentrifugation. The control experiment using BM+0.5% BSA showed no fluorescently labelled particles (data not shown).

#### Cell migration assay with different concentration of isolated EVs

The actual biological effects of different EVs isolates were investigated using the scratch wound assay, which was modified to measure the wound closure of cells towards the “wound”. Lung epithelial cells (BEAS-2B) was selected and 10 μg/mL Lipopolysaccharides (LPS) was used before wound scratch as an *in vitro* model of acute lung injury. LPS is a key pathogenic factor that induces various inflammatory mediators, which resulted in lung inflammatory and epithelial damage [51]. Concentrations of CEVs and DEVs isolated using TFF and ultracentrifugation, ranging from 10 to 1000 EVs per cell, were used in order to test their ability to promote cell migration and increase wound closure after injury. Both CEV and DEV treated cells enhanced wound closure percentage when compared with LPS-treated only cells **(Fig. 6)**. Increasing the concentration of CEVs isolated using both TFF and ultracentrifugation (from 10 to 1000 EVs per cell) increased the wound closure percentage of BEAS-2B cells **(Fig. 6 A, C)**. Especially, at the concentration of 10 EVs per cell for CEVs isolated using TFF, the wound closure percentage was significantly increased as compared to LPS-treated only cells **(Fig. 6 A)**.

**Fig. 6.**
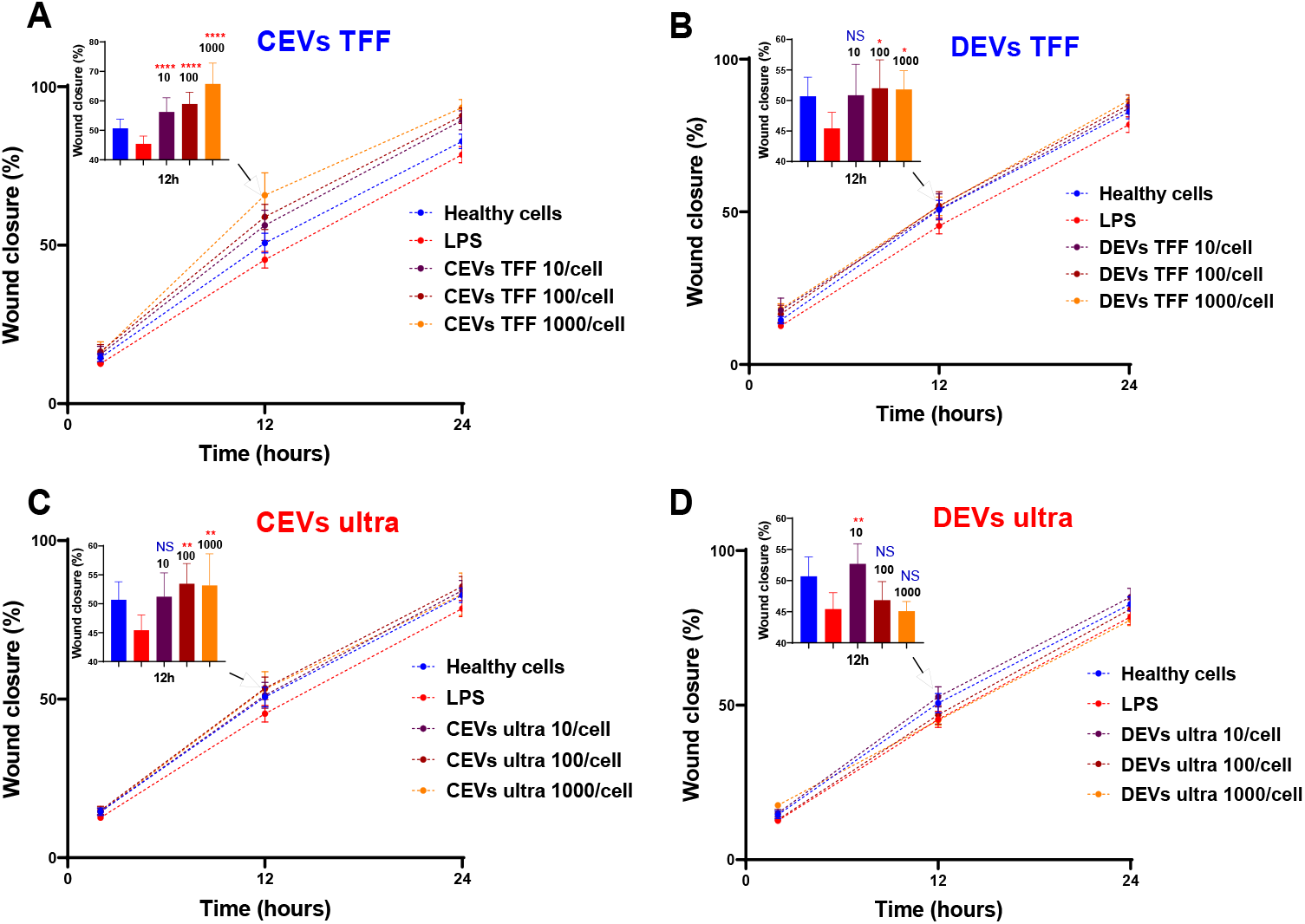
Assessment of wound closure in injured BEAS-2B in response to EV treatment. BEAS-2B cells were ‘injured’ with 10 μg/mL LPS and treated with CEVs isolated using **(A)** TFF and **(C)** ultracentrifugation; DEVs isolated using **(B)** TFF and **(D)** ultracentrifugation. Error bars represent standard deviation (SD), n = 8. Statistics significance was determined by ordinary one-way ANOVA test, followed by Dunn’s multiple comparison’s test for pair-wise comparisons. Statistical significance shown were between LPS treatment and other groups at time point 12 h. *: *P* < 0.1; **: *P* < 0.01; ****: *P* < 0.0001; NS: not significant.

Treatment with DEVs isolated using TFF resulted in increased wound closure percentage of BEAS-2B cells **(Fig. 6 B)**. The wound closure percentage of BEAS-2B was significantly increased at the concentration of 100 and 1000 EVs per cell for DEVs isolated using TFF. In contrast, with high concentration of DEVs isolated using ultracentrifugation (100 and 1000 EVs per cell), the wound closure percentage remain as low as the LPS-treated cells **(Fig. 6 D)**. This was supported by the highest sedimented amount for DEVs isolated using ultracentrifugation using the DG model, which resulted in the highest dose for cells. We postulate that high DEVs dose could be causing adverse effects on the cells.

#### Cellular stress after injury in response to different concentration of isolated EVs

Nitric oxide (NO) measurement allows the measurement of cellular stress levels in response to injury [52]. NO participates in diverse physiological and pathological processes, such as inflammation [53]. In response to inflammatory stimuli, NO production is markedly elevated [54]. Given the importance in studying understanding NO for health and disease, a number of fluorescent sensors have been reported to date [55]. However, a common feature of small molecule sensors is poor water solubility, arising from the highly aromatic structures. This has two main drawbacks in cellular studies: firstly, a need to prepare a stock solution of the dye in an organic solvent such as dimethyl sulfoxide (DMSO), which itself can perturb the system; and secondly, a tendency for the dye to aggregate and self-quench. We therefore sought to prepare a water soluble fluorescent sensor for NO by utilising the previously-reported selective conversion of aromatic *ortho*-diamines to triazoles in the presence of nitric oxide and oxygen [55], conjugating this reactive group to a 4-amino-1,8-naphthalimide fluorophore, which we have previously shown to have great potential for sensing applications [56, 57]. Water solubility was achieved by incorporating a triethylene glycol (TEG) group to give the final probe, NpNO1 **(Fig. 7 A)**. NpNO1 was prepared in 4 synthetic steps from commercially-available bromoacetic naphthalic anhydride, in 42% overall yield as detailed in the Supplementary Information **(Scheme S1)**. NpNO1 was found to have excellent aqueous solubility up to 1 mM **(Fig. S6)** and a strong fluorescence turn-on at 455 nm with NO addition **(Fig. 7 B)**.

**Fig. 7.**
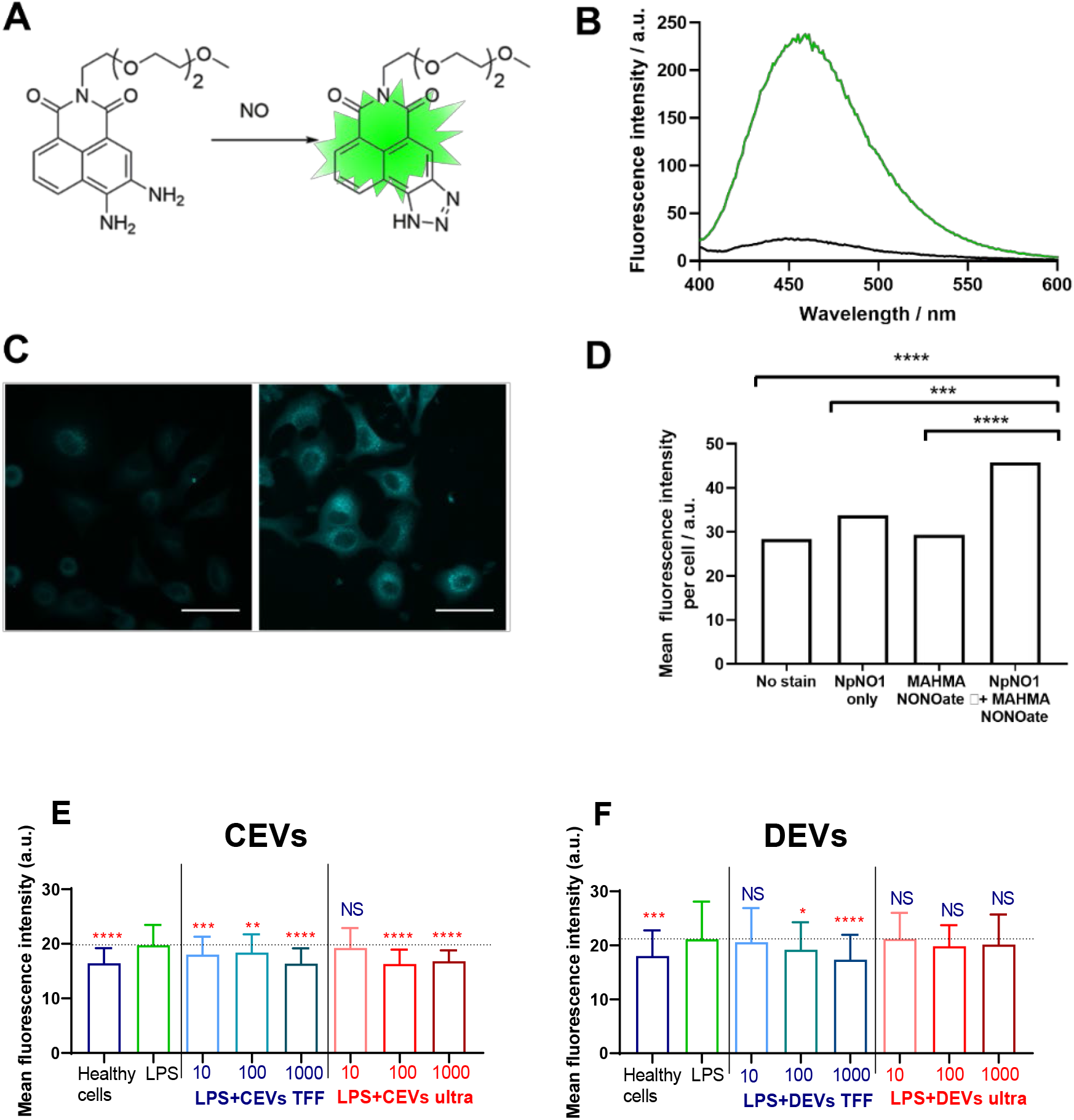
A novel, water soluble fluorescent sensor for intracellular NO and assessing NO levels in BEAS-2B cells. **(A)** Water-soluble NO sensor NpNO1 and the product of its reaction with nitric oxide to give a fluorescent triazole derivative. **(B)** Fluorescence emission response of NpNO1 (10 μM) in the presence of diethylamine NONOate (1 mM). **(C)** Representative micrographs of A549 cells treated with NpNO1 (50 μM) in the absence (left) and presence (right) of MAHMA NONOate (5 mM). Scale bar represents 50 μm. **(D)** Mean fluorescence intensity of A549 cells treated as indicated measured across at least 3 different field of views in each of the 3 independent experiments performed (n = 3). MAHMA NONOate = (Z)-1-[N-Methyl-N-[6-(N-methylammoniohexyl)amino]]diazen-1-ium-1,2-diolate, an NO donor. BEAS-2B cells were ‘injured’ with 10 μg/mL LPS and dosed with **(E)** CEVs isolated using TFF and ultracentrifugation and **(F)** DEVs isolated using TFF and ultracentrifugation. Statistics significance was determined by ordinary one-way ANOVA, followed by Dunn’s multiple comparison’s test for pair-wise comparisons, n = 3. Statistical significance shown were between LPS treatment and other groups. *: *P* < 0.1; **: *P* < 0.01; ***: *P* < 0.001; ****: *P* < 0.0001; NS: not significant. Error bars denote standard deviation (SD). The dash line represents for the mean level of LPS group.

The ability of NpNO1 to detect changes in intracellular NO levels by confocal microscopy was confirmed in A549 (human alveolar basal epithelial) cells treated with 50 μM of NpNO1 overnight, in the presence or absence of 5 mM MAHMA NONOate (**Fig. 7 C, Fig. S6)**. MAHMA NONOate spontaneously releases NO, and hereafter will be referred to as the NO donor. We observed minimal/basal fluorescence from untreated cells (no stain), cells treated with only MAHMA NONOate or cells treated with NpNO1 alone. In A549 cells treated with both the NpNO1 and the NO donor, we observed fluorescent puncta in every cell, and a significant increase in mean fluorescence intensity.

We then assessed intracellular NO levels after adding various EVs, by quantifying the NpNO1 fluorescence intensity in cells imaged using confocal microscopy **(Fig. S7)**.

Similar to the scratch wound assay, we modelled cellular injury by treating human BEAS-2B epithelial cells with 10 μg/mL LPS. The intracellular NO, as indicated by the fluorescence intensity of NpNO1, increases after LPS injury **(Fig. 7)**, which is consistent with the literature [58]. Next, we assessed the fluorescence intensity of NpNO1, which corresponds to the levels of intracellular NO present, in LPS-treated cells dosed with various isolated EVs. Upon adding the concentration of 10-1000 EVs/cells for CEVs isolated using TFF, the NO levels diminished as compared to LPS-treated cells **(Fig. 7 E)**. Whereas, dosing the LPS-treated cells with 10 EVs/cell for CEVs isolated using ultracentrifugation, the intracellular NO levels remain high, as in LPS-treated cells **(Fig. 7 E)**. The intracellular NO levels decreased when 100 and 1000 EVs/cells for CEVs isolated using ultracentrifugation were added compared to LPS-treated cells.

The NpNO1 fluorescence intensity remained as high as LPS-treated cells using the concentration of 10 DEVs/cell isolated using TFF **(Fig. 7 F)**. The intracellular NO levels, as indicated by NpNO1 fluorescence intensity, decreased when the cells were treated with 100 and 1000 DEVs isolated using TFF. For DEVs isolated using ultracentrifugation, the intracellular NO levels remain as high as the LPS-treated cells at all concentrations **(Fig. 7 F)**.

## Discussion

While there is palpable excitement about the future of EV-based medicine, a current major limitation is the lack of understanding about how to effectively and uniformly isolate them from complex biological milieu [21]. Furthermore, it has not been clear whether different isolation methods extract different sub-populations of EVs, which would impact on their downstream applications. This gap in knowledge is associated with conceptual and technical limitations in EV characterisation. We have addressed this challenge by using multiple techniques including optical, non-optical and high-resolution single vesicle characterisation methods, and we provide extensive experimental evidence that *isolation method determines the composition and biological function of EV isolates.*

Two common isolation methods were used in this study: ultracentrifugation and TFF. Using four different techniques, we demonstrated that the size distributions of EVs isolated using ultracentrifugation and TFF were different. Since EVs are heterogenous and each characterisation technique has its own limits of detection [32], it is essential to measure the size distribution using multiple techniques. In addition, for the first time we have demonstrated that large EVs (>100 nm) isolated using ultracentrifugation have a higher dry mass than large EVs isolated using TFF. Here we have introduced the parameter of EV dry mass as a metric to reveal the total mass of molecular cargo inside each vesicle, a measurement not previously made in the EV field. The differences in dry mass between EV isolates suggested different molecular composition. Indeed, we showed that there were variations between the expression levels of surface markers (CD9, CD63 and CD81) between different EV isolates at sub-population level by using nano flow cytometry. It has been reported that the presence of each surface maker in EV populations was different depending on the isolation methods of EVs [59], which clearly indicated that different subpopulations of EVs have been extracted using different isolation methods. Furthermore, our study showed that the amount of nucleic acid content in EVs isolated using TFF was higher than in EVs isolated using ultracentrifugation for both EV types. The higher nucleic acid content for EVs isolated using TFF was further confirmed at individual vesicle characterisation using atomic force nanoscale infrared spectroscopy. By combining multiscale characterisation techniques, our study allows robust and precise quantification of molecular composition of EVs. This methodology goes a step beyond conventional characterisation methods such as immunoassays or mass spectroscopy, which not only lack single vesicle and subpopulation resolution and flexibility in practical applications but also are very expensive [60]. This study paves the way to the future diagnostic and therapeutic applications of EVs that depend on identification/quantification of specific cargo.

We have also showed for the first time that the variations in the physicochemical properties of different EV types correlate to their different interactions with cell membranes. By using a distorted grid model to predict how quickly different types of EVs can reach the cell membrane, we demonstrated that the predicted sedimented amount of EVs isolated using ultracentrifugation were 60-100 times higher than EVs isolated using TFF. The higher sedimentation of EVs isolated using ultracentrifugation could be due to the presence of agglomerates, which is consistent with the size distribution and mass results. The agglomerates in EVs isolated using ultracentrifugation was confirmed further using correlative holotomography and fluorescence microscopy. By using sedimentation and cellular uptake of EVs, the actual cellular response that mediated by EV uptake are elucidated. We showed that the higher sedimentation of EVs influenced their effects on cell migration and cellular stress post-injury. DEVs isolated using ultracentrifugation with highest predicted sedimentation showed negative effects on both cell migration and cellular stress post-injury. This finding emphasizes that EV agglomerates, which sedimented faster, affect the biological response in cells. Differences in biological responses to the EV isolates were confirmed using newly developed nitric oxide (NO) probe, which allowed for quantitative and qualitative assessment of intracellular stress [52]. NO probe enabled to determine the ability of each EV isolates to promote cell recovery from injury.

In conclusion, we have shown that the physicochemical properties, molecular composition, the presence of surface markers and subsequent biological effects of EVs isolated using ultracentrifugation are markedly different from those isolated by TFF. This study confirms that the isolation method determines which the composition of EV sub-populations isolated. Demonstrated here correlation between physicochemical properties and biological effects of EVs, confirmed that the downstream applications of EVs are determined by the effectiveness of isolation methods to isolate and fractionate different subpopulations of EVs.

The methodology that we present here for the high resolution and multiscale measurement of physicochemical and functional properties of EVs is likely to accelerate progress in the development and refinement of isolation methods. Our findings, which uncovered that different EV subpopulations are isolated by different methods, shed new light on our understanding of EV secretion by cells. Furthermore, knowing what the differences in EV composition are depending on the isolation method will support the development of accurate EV-based diagnostic tools for early disease detection. We will also be able to define populations of EVs which have therapeutic potential for specific medical conditions. Taken together, presented study advanced current understanding of the effect of isolation methods on EV composition and functionality. Our results are likely to contribute to future EV research and provide a backbone for rapid translation of EVs to practical applications.

## Materials and methods

### Cell culture and maintenance

Both chorionic and decidual MSCs cell lines (CMSC29 and DMSC23) were obtained from Royal Women’s Hospital in Melbourne, Australia. CMSC29 cells were cultured in 85% AmnioMAX™ C-100 basal medium and 15% AminoMAX™ C-100 supplement (Invitrogen™, ThermoFisher Scientific). DMSC23 cells were cultured in MesenCult™ MSC Basal medium (Human), 10% Mesenchymal stem cell stimulatory supplement (STEMCELL Technologies, Canada), GlutaMAX^TM^ (Life Technologies, Australia) and antibiotics (Pen/Strep) (100 units penicillin and 0.1 mg/mL streptomycin, Sigma-Aldrich, Australia). BEAS-2B cells were cultured in medium containing Dulbecco’s Modified Eagle’s Medium (DMEM medium-high glucose, Sigma-Aldrich, Australia) supplemented with 10% Foetal Bovine Serum (FBS, Bovogen, Australia) and antibiotics (Pen/Strep). Cells were subcultured every 2-3 days and maintained in the incubator at 37°C supplemented with 5% CO_2_. hanks’ balanced salt solution (HBSS, Sigma-Aldrich, Australia) was used for washing CMSC29 and DMSC23 cells. TrypLE™ Express (Gibco, Denmark) was used to dissociate the adherent cells. EV isolation and collection were from CMSC29 and DMSC23 cells at passages P23-28.

### Isolation of EVs by ultracentrifugation

CMSC29 and DMSC23 cells were cultured to 80% confluency. Cells were washed twice with HBSS before incubating cells with EV isolation media (MesenCult™ MSC Basal medium containing 0.5% (w/v) bovine serum albumin (BSA, Sigma-Aldrich, Australia) for 48 h. After 48 h, EV-containing media was collected and centrifuged at 500 × g for 5 min and 2,000 × g for 10 min to remove cells and debris. The supernatant was then transferred to thick-wall polycarbonate ultracentrifuge tubes (Seton Scientific Inc, USA) and centrifuged at 100 000 × g for 60 min at 4°C using rotor Ti-70 in an Optima LE-80K Ultra Centrifuge (Beckman Coulter, Australia). The harvested EV pellet was resuspended in 1 mL RNase-free Phosphate buffered saline (RNase-free PBS, Lonza, Australia) and washed using ultracentrifugation at 100,000 × g for 60 min at 4°C. The pellet was resuspended in 1 mL RNase-free PBS and transferred to RNase-free microcentrifuge tubes. The EV pellets were stored at 4°C to avoid losing biological function during the freezing process.

### Isolation of EVs by tangential flow filtration (TFF)

After removing cells and debris by centrifugation 500 × g (5 min) and 2000 × g (10 min) as described above, the EV containing supernatant was filtered (0.45 μm) and transferred to TFF-Easy 20 kDa MWCO (HansaBioMed/Lonza, Tallinn, Estonia) for EV concentration. The EVs concentration process was described in the manufacture’s protocol (HansaBioMed/Lonza) and EVs were finally diafiltrated in RNase-free PBS.

### Size and concentration measurement using Nano-flow cytometry (nFCM)

The size and concentration of EVs were measured using NanoFCM (Xiamen Fuliu Biological Technology Co., Ltd, Xiamen, China). A mixture of silica nanospheres (68, 91, 113 and 155 nm) was used as the size standard for the construction of a calibration curve and standard 200 nm polystyrene spheres were used for laser alignment. All events were collected for 120 s and size (SSC) triggering was used to detect EVs. The total events collected ranging from 3000-6000 events.

### Size and concentration measurement using Particle Tracking Analysis (PTA)

EV samples were diluted with RNase free water to achieve concentration between 1 × 10^8^ and 1 × 10^9^ EVs/mL and measured using a NanoSight NS300 (Malvern Panalytical Ltd, Malvern, United Kingdom). A syringe pump with speed 40μl/mL and cell temperature was set at 25°C. Embedded laser wavelength was 488 nm and the particles were imaged with an auto-focus camera for 60s. Data were obtained at camera-level 11. The analysis settings were set to “auto”, and the detection threshold was set to 5 in the NanoSight Software NTA (version 3.2) to assess mean and modal particle diameters, D50 values (which represents the 50^th^ percentile of the averaged cumulative number-weighted particle size distribution) and particle number concentration. Triplicate measurements were made for each EV sample.

### Size measurement using Dynamic Light Scattering (DLS)

EV samples were diluted in RNase-free water to achieve a particle concentration ranging from 1 × 10^9^ to 1 × 10^10^ EVs/mL and were measured in a Zetasizer Ultra (Malvern Panalytical Ltd, Malvern, United Kingdom). The manufacture’s default software setting for EVs (liposomes) was selected and three cycles were performed for each measurement at 4°C. Data were analysed using general purpose mode in ZS XPLORER software (version 1.2.0.91) and the size distribution of EV populations was presented as a percentage of intensity.

### Size and concentration measurement using Tunable Resistive Pulse Sensing (TRPS)

TRPS was performed using a qNano (IZON, New Zealand) to measure the particle size and the concentration of EVs. EVs were suspended in electrolytes and passed through an engineered pore (NP100), which provided direct measurement of size and concentration. Buffers and reagents were freshly prepared and filtered (0.22 μm) before the measurement. The detailed protocol for qNano measurements was described in Vogel *et al.* study [61].

### Morphology analysis of EVs using Atomic Force Microscopy (AFM)

EV samples were placed onto zinc selenide prism, dried overnight and subsequently imaged using atomic force microscopy (AnasysInstruments, USA). Images were obtained in contact mode at a scan rate of 0.5 Hz using EX-T125 probe with nominal resonance frequency 200-400 kHz and spring constant 13-77 Nm^-1^ (AnasysInstruments, USA).

### Dry mass measurement of EVs using Resonant mass measurement (RMM)

The buoyant mass of the particles was measured with an Archimedes Particle Metrology System (Malvern Panalytical, Malvern, United Kingdom). The microchannel sensor used consisted of a microfluidic channel with a cross section of 2 × 2 μm^2^. The measured buoyant mass was used to determine the size and dry mass of the EVs, assuming an EV density of 1.4 g/cm^3^ (calculated from measurements of EVs of known diameter, nominally 200 nm **(Supplementary Fig. S1)**, known fluid densities and assuming a spherical model. The sensitivity factor of the microchannel resonator was determined using monomodal gold calibration particles (NIST RM 8016, 60 nm, USA). The limit of detection (LOD) for dry mass is 10^-15^ g and for size distribution is 100 nm.

### Quantification of EV surface markers using nFCM

The monoclonal antibodies, anti-CD9 Alexa Fluor^®^ 488-conjugated (R & D systems, Canada) or anti-CD63 Alexa Fluor^®^ 488-conjugated (Invitrogen, ThermoFisher Scientific) or anti-CD81 Alexa Fluor^®^ 488-conjugated (R & D systems, Canada) were used to assess protein surface markers of EVs. Approximately 1 × 10^10^ EVs/mL was stained using 8 μg/mL of each antibody and incubated 30 min at 37°C in dark condition. The EV samples were then washed with 2 mL RNase free PBS for three times using ultracentrifugation 100 000 × g, 4°C, 70 min each. The supernatants were carefully aspirated from the bottom of the tubes in every wash. Subsequently, the pellets were dissolved in RNase free PBS and the fluorescence intensity was measured using a nFCM. The control sample was the basal medium with 0.05% (w/v) BSA without EVs. All fluorescence events were detected in FITC triggering of nFCM and the threshold level was set by default in NF Profession 1.0 acquisition software.

### Quantification of nucleic acid using nFCM

EVs isolated using TFF and ultracentrifugation were stained using 10 μM SYTO RNASelect green fluorescent cell stain (Invitrogen™, ThermoFisher Scientific) at 37°C for 30 min. The EV samples were loaded on the exosome spin column MW 3000 (Invitrogen™, ThermoFisher Scientific) to remove unbound dyes and measured using nFCM. The control sample was basal medium without EVs. All fluorescence events were collected for 120 s and fluorescence (FITC) triggering of a nFCM was used to detect fluorescence EVs. Data were analysed using FlowJo software (version 10.6). The threshold level was set above the background level by the NF Profession 1.0 acquisition software by default.

### Quantification of lipid content using nFCM

PKH67 and Diluent C (ThermoFisher Scientific) was selected to be the general membrane labelling for EVs. PKH67 was diluted in 100 μL Diluent C to a final concentration of 15 μM (dye solution). Approximately 1 × 10^10^ EVs/mL were diluted with 80 μL Diluent C, added to dye solution, and incubated for 3 min with gentle pipetting. Excess dye was bound with 10% (w/v) BSA in RNase free water. The EV samples were then diluted to 2 mL with RNase free PBS and washed three times using ultracentrifugation 100 000 × g, 4°C, 70 min each. The pellet was gently resuspended in 100 μL RNase free PBS and nFCM was used to measure the fluorescence intensity of EVs. The control sample were the basal medium only and the basal medium with 0.05% (w/v) BSA without EVs. All fluorescence events were triggered in the FITC channel and collected for 120 s. The threshold was set by default in NF Profession 1.0 acquisition software.

### Molecular composition analysis of EVs using atomic force nanoscale infrared spectroscopy (AFM-IR)

The protocol for EV characterisation using AFM-IR (nanoIR, AnasysInstruments, USA) was described in our previous study [62]. Briefly, each EV sample was placed on a zinc selenide prism and dried overnight. The laser signal was optimized before acquiring the nanoIR spectra ranging from 1000-1800 cm^-1^ at 4 cm^-1^ intervals with a scan rate of 0.5 Hz. A gold coated tip and a silicon nitride cantilever with a nominal spring constant of 0.5 Nm^-1^ were used for all measurements. The acquired scan sizes were 10 × 5 μm for each sample and Analysis Studio™ software was used for data analyses. The ‘Savitzky-Golay’ function was used to achieve smoothing of the spectra with the polynomial function of 3, and 8 numbers of points.

### Measurement of predicted sedimentation of EVs – distorted grid (DG) model

The predicted transport modelling of TFF and ultracentrifugation isolated EVs, which was originally applied for the engineered nanomaterials (ENMs), was modified and adapted for EVs [63]. Briefly, the protocol comprised three interconnected parts: ENM dispersion preparation and characterisation in suspension, effective density calculation, and delivered dose computation [63]. The dispersion preparation and characterisation were not applied to EVs. The delivered dose metric was used to calculate the fraction of administered EVs in 96 well plate over 24 h. Data were acquired using Matlab.

The effective density of EVs (*Peffective density*) was determined using the equation:

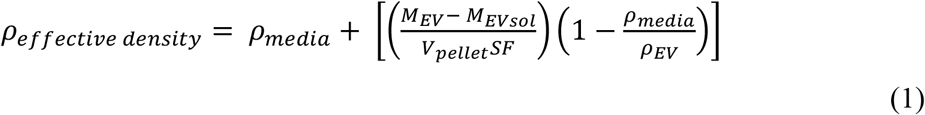

whereby:

*ρ_media_* is the density of medium (g/cm^3^).
*M_EV_* is the total mass of EVs (g) in the dispensed volume of suspension.
*M_EVsoi_* is the mass of dissolved EVs in the dispensed volume of suspension.
*V_pellet_* is the measured pellet size in centimeters squared inside PCV tube.
*SF* is the stacking factor, which is the potion of the pellet that is composed of agglomerates (theoretical maximum 0.74 for ordered stacking).
*ρ_EV_* is the density of EVs (g/cm^3^).

### Visualization of EV uptake using holotomography and fluorescence microscopy

CMSC29 EVs isolated using TFF and ultracentrifugation were labelled with PKH67 followed the lipid staining protocol as described in section 2.7.4. The staining of EVs was done 1 h prior to incubation. Approximately 2 × 10^4^ BEAS-2B cells were dosed with 1 × 10^9^ PKH67-stained EVs/mL and incubated with the dye for 3 h. Uptake of EVs by cells was performed on a holotomography microscope Tomocube HT-2H (Tomocube Inc., Daejeon, Korea). A water immersion objective (60 ×, N.A = 1.2) was used to acquire the images. Z-stacked images were acquired across each field-of-view, with a minimum of 4 field-of-views imaged for each type of EVs. The images were acquired in TomoStudio™ 2.0 software and they were further analysed using ImageJ FIJI.

### Analysis of cell migration to EV treatment after LPS injury

Cell migration in the presence of isolated EVs was measured by comparing the area of closure of a two-dimensional scratch wound. BEAS-2B cells were seeded at 1 × 10^4^ cells per well on Image Lock 96-well plates and allowed to adhere overnight. ‘Injury’ was induced using 10 μg/mL lipopolysaccharides (LPS) for 24 h [64]. A wound on the midline of culture well was then created using a 96-pin wound making tool (IncuCyte^^®^^ WoundMaker^TM^). After washing the cells with RNase free PBS once, EVs isolated using TFF and ultracentrifugation were added at ascending concentration ranging from 10-1000 EVs per cell. Wound images were taken every 2 h with a 10 × magnification objective lens using the IncuCyte^®^ live cell imaging system and IncuCyte ZOOM software program (Essen BioScience, USA). Wound confluence (%), which was represented as the wound closure (%), was assessed for all images using IncuCyte ZOOM software. Data were analysed using GraphPad and measurements of 8 samples (n=8) were performed for each condition.

### Analysis of cellular responses to EV treatment after LPS injury

Cellular responses post-injury in the presence of TFF and ultracentrifugation isolated EVs were assessed by measuring intracellular nitric oxide (NO) levels. Approximately 5 × 10^3^ BEAS-2B cells were seeded on glass bottom dish precoated in L-poline (MatTek) and allowed to adhere overnight. Cells were then exposed to 10 μg/mL of LPS for 24 h. Cells were next incubated with 50 μM NpNO1 probe for 24 h at 37°C, 5% CO_2_. Excess NpNO1 probe was washed and imaged in FluoroBrite™ DMEM media (Gibco, ThermoFisher Scientific) supplemented with 10% FBS and antibiotics (Pen/Strep). Samples were imaged using Olympus FV3000 microscope equipped with a 405 nm laser, a water 60× objective lens and an incubator stage maintained at 37°C, and 5% CO_2_. A minimum of 3 field-of-views were imaged for each condition per experiment with Z-stacked images per field-of-view. Maximum projected micrographs of the Z-stacks were presented in this study. Regions of interest were drawn around each cell and mean fluorescence intensities were quantified using ImageJ FIJI. Statistical analyses were performed on the mean fluorescence intensity values and plotted using GraphPad.

### Statistical analyses

Data were analysed and presented as mean ± standard deviation (SD). Ordinary one-way ANOVA followed by Dunn’s multiple comparison’s test for pair-wise comparisons was used to determine the differences between multiple groups. A *P*-value < 0.05 is considered to be statistically significant.

## Supporting information

Supplementary data including graphical abstract

## List of Abbreviations

AFM: Atomic Force Microscopy;
AFM-IR: Atomic Force Microscope InfraredSpectroscopy;
CMSC29: chorionic mesenchymal stromal/stem cell line;
CEVs: extracellular vesicles derived from chorionic mesenchymal stromal/stem cell;
DMSC23: decidual mesenchymal stromal/stem cell line;
DEVs: extracellular vesicles derived from decidual mesenchymal stromal/stem cell;
EVs: extracellular vesicles;
HBSS(-): hanks’ balanced salt solution;
LPS: lipopolysaccharide;
MSC: mesenchymal stromal/stem cell;
PTA: Particle Tracking Analysis;
nFCM: Nano-flow Cytometry;
RMM: Resonant Mass Measurement;
TRPS: Tunable Resistive Pulse Sensing;
TFF: tangential flow filtration.

## Acknowledgements

The authors acknowledge the facilities and the scientific and technical assistance of the Bosch Molecular Biological Facility and Australian Microscopy & Microanalysis Research Facility at the Australian Centre for Microscopy & Microanalysis, The University of Sydney, Tomocube company, Republic of Korea.

The authors would like to acknowledge the Australian Research Council (DP180101353, DP180101897) for funding, the Westpac Scholars Trust for a Research Fellowship (EJN), the University of Sydney for a SOAR Fellowship (EJN, WCh), the Australian government for Research Training Program Scholarships (KGL), and Medical Advances Without Animals Trust MAWA (THP, WCh).

## Declaration of Interest Statement

The authors have declared no conflict of interest.

## Notes

### Competing Interest Statement

The authors have declared no competing interest.

### Summary of Updates

Tittle revised, author name updated

## References

1. Iraci, N., et al., Focus on Extracellular Vesicles: Physiological Role and Signalling Properties of Extracellular Membrane Vesicles. International journal of molecular sciences, 2016. 17(2): p. 171–171.

2. Lötvall, J., et al., Minimal experimental requirements for definition of extracellular vesicles and their functions: a position statement from the International Society for Extracellular Vesicles. Journal of extracellular vesicles, 2014. 3: p. 26913–26913.

3. Candelario, K.M. and D.A. Steindler, The role of extracellular vesicles in the progression of neurodegenerative disease and cancer. Trends in molecular medicine, 2014. 20(7): p. 368–374.

4. Taylor, D.D. and C. Gercel-Taylor, MicroRNA signatures of tumor-derived exosomes as diagnostic biomarkers of ovarian cancer. Gynecol Oncol, 2008. 110(1): p. 13–21.

5. Danielson, K.M. and S. Das, Extracellular Vesicles in Heart Disease: Excitement for the Future ? Exosomes and microvesicles, 2014. 2(1): p. 10.5772/58390.

6. Yanez-Mo, M., et al., Biological properties of extracellular vesicles and their physiological functions. Journal of Extracellular Vesicles, 2015. 4.

7. György, B., et al., Therapeutic applications of extracellular vesicles: clinical promise and open questions. Annual review of pharmacology and toxicology, 2015. 55: p. 439–464.

8. Piffoux, M., et al., Imaging and Therapeutic Potential of Extracellular Vesicles, in Design and Applications of Nanoparticles in Biomedical Imaging, J.W.M. Bulte and M.M.J. Modo, Editors. 2017, Springer International Publishing: Cham. p. 43–68.

9. Yekula, A., et al., From laboratory to clinic: Translation of extracellular vesicle based cancer biomarkers. Methods, 2020. 177: p. 58–66.

10. Ramirez, M.I., et al., Technical challenges of working with extracellular vesicles. Nanoscale, 2018. 10(3): p. 881–906.

11. Simonsen, J.B. and R. Munter, Pay Attention to Biological Nanoparticles when Studying the Protein Corona on Nanomedicines. Angewandte Chemie-International Edition, 2020. 59(31): p. 12584–12588.

12. Sunkara, V., H.K. Woo, and Y.K. Cho, Emerging techniques in the isolation and characterization of extracellular vesicles and their roles in cancer diagnostics and prognostics. Analyst, 2016. 141(2): p. 371–381.

13. Gardner, L., et al., The biomolecule corona of lipid nanoparticles contains circulating cell-free DNA. Nanoscale Horiz, 2020. 5(11): p. 1476–1486.

14. Mateescu, B., et al., Obstacles and opportunities in the functional analysis of extracellular vesicle RNA – an ISEV position paper. J Extracell Vesicles, 2017. 6(1): p. 1286095.

15. Thery, C., et al., Minimal information for studies of extracellular vesicles 2018 (MISEV2018): a position statement of the International Society for Extracellular Vesicles and update of the MISEV2014 guidelines. J Extracell Vesicles, 2018. 7(1): p. 1535750.

16. Gardiner, C., et al., Techniques used for the isolation and characterization of extracellular vesicles: results of a worldwide survey. Journal of extracellular vesicles,2016.5: p. 32945–32945.

17. Ismail, N., et al., Macrophage microvesicles induce macrophage differentiation and miR-223 transfer. Blood, 2013. 121(6): p. 984–995.

18. Kang, H., J. Kim, and J. Park, Methods to isolate extracellular vesicles for diagnosis. Micro and Nano Systems Letters, 2017. 5(1): p. 15.

19. Witwer, K.W., et al., Standardization of sample collection, isolation and analysis methods in extracellular vesicle research. J Extracell Vesicles, 2013. 2.

20. Coumans, F.A.W., et al., Methodological Guidelines to Study Extracellular Vesicles. Circ Res, 2017. 120(10): p. 1632–1648.

21. Li, X., et al., Challenges and opportunities in exosome research-Perspectives from biology, engineering, and cancer therapy. APL Bioeng, 2019. 3(1): p. 011503.

22. Momen-Heravi, F., et al., Current methods for the isolation of extracellular vesicles. Biol Chem, 2013. 394(10): p. 1253–62.

23. Busatto, S., et al., Tangential Flow Filtration for Highly Efficient Concentration of Extracellular Vesicles from Large Volumes of Fluid. Cells, 2018. 7(12).

24. Heath, N., et al., Rapid isolation and enrichment of extracellular vesicle preparations using anion exchange chromatography. Scientific Reports, 2018. 8(1): p. 5730.

25. Kim, S.Y., et al., Placenta Stem/Stromal Cell Derived Extracellular Vesicles for Potential Use in Lung Repair. PROTEOMICS, 2019. 19(17): p. 1800166.

26. Qin, S.Q., et al., Establishment and characterization of fetal and maternal mesenchymal stem/stromal cell lines from the human term placenta. Placenta, 2016. 39: p. 134–46.

27. Bishop, J.B., J.C. Martin, and W.M. Rosenblum, A Light-Scattering Method for Qualitatively Monitoring Aggregation Rates in Macromolecular Systems. Journal of Crystal Growth, 1991. 110(1-2): p. 164–170.

28. Barnett, C.E., Some applications of wave-length turbidimetry in the infrared. Journal of Physical Chemistry, 1942. 46(1): p. 69–75.

29. Gercel-Taylor, C., et al., Nanoparticle analysis of circulating cell-derived vesicles in ovarian cancer patients. Anal Biochem, 2012. 428(1): p. 44–53.

30. Tosi, M.M., et al., Chapter Six – Dynamic light scattering (DLS) of nanoencapsulated food ingredients, in Characterization of Nanoencapsulated Food Ingredients, S.M. Jafari, Editor. 2020, Academic Press. p. 191–211.

31. Varga, Z., et al., Size Measurement of Extracellular Vesicles and Synthetic Liposomes: The Impact of the Hydration Shell and the Protein Corona. Colloids Surf B Biointerfaces, 2020. 192: p. 111053.

32. Gandham, S., et al., Technologies and Standardization in Research on Extracellular Vesicles. Trends Biotechnol, 2020.

33. Maas, S.L.N., et al., Possibilities and limitations of current technologies for quantification of biological extracellular vesicles and synthetic mimics. Journal of Controlled Release, 2015. 200: p. 87–96.

34. Cohen, J., et al., Interactions of engineered nanomaterials in physiological media and implications for in vitro dosimetry. Nanotoxicology, 2013. 7(4): p. 417–31.

35. Rupert, D.L.M., et al., Methods for the physical characterization and quantification of extracellular vesicles in biological samples. Biochim Biophys Acta Gen Subj, 2017. 1861(1 Pt A): p. 3164–3179.

36. Linares, R., et al., High-speed centrifugation induces aggregation of extracellular vesicles. Journal of Extracellular Vesicles, 2015. 4(1): p. 29509.

37. Taylor, D.D. and S. Shah, Methods of isolating extracellular vesicles impact downstream analyses of their cargoes. Methods, 2015. 87: p. 3–10.

38. Nordin, J.Z., et al., Ultrafiltration with size-exclusion liquid chromatography for high yield isolation of extracellular vesicles preserving intact biophysical and functional properties. Nanomedicine, 2015. 11(4): p. 879–83.

39. Ohno, S., et al., Systemically injected exosomes targeted to EGFR deliver antitumor microRNA to breast cancer cells. Mol Ther, 2013. 21(1): p. 185–91.

40. Lai, C.P., et al. Visualization and tracking of tumour extracellular vesicle delivery and RNA translation using multiplexed reporters. Nature communications, 2015. 6, 7029 DOI: 10.1038/ncomms8029.

41. Takov, K., D.M. Yellon, and S.M. Davidson Confounding factors in vesicle uptake studies using fluorescent lipophilic membrane dyes. Journal of extracellular vesicles, 2017. 6, 1388731 DOI: 10.1080/20013078.2017.1388731.

42. Kim, S.Y., et al., None of us is the same as all of us: resolving the heterogeneity of extracellular vesicles using single-vesicle, nanoscale characterization with resonance enhanced atomic force microscope infrared spectroscopy (AFM-IR). Nanoscale Horizons, 2018. 3(4): p. 430–438.

43. Kim, S.Y., et al., High-fidelity probing of the structure and heterogeneity of extracellular vesicles by resonance-enhanced atomic force microscopy infrared spectroscopy. Nature Protocols, 2019. 14(2): p. 576–593.

44. Lee, J., et al., Infrared spectroscopic characterization of monocytic microvesicles (microparticles) released upon lipopolysaccharide stimulation. Faseb j, 2017. 31(7): p. 2817–2827.

45. Mohan, V., et al., Hydroxide-catalyzed cleavage of selective ester bonds in phosphatidylcholine: An FTIR study. Vibrational Spectroscopy, 2020. 109.

46. Balan, V., et al., Vibrational Spectroscopy Fingerprinting in Medicine: from Molecular to Clinical Practice. Materials (Basel), 2019. 12(18).

47. Willms, E., et al., Extracellular Vesicle Heterogeneity: Subpopulations, Isolation Techniques, and Diverse Functions in Cancer Progression. Front Immunol, 2018. 9: p. 738.

48. DeLoid, G.M., et al., Advanced computational modeling for in vitro nanomaterial dosimetry. Part Fibre Toxicol, 2015. 12: p. 32.

49. DeLoid, G.M., et al., Preparation, characterization, and in vitro dosimetry of dispersed, engineered nanomaterials. Nature Protocols, 2017. 12(2): p. 355–371.

50. Kim, S., et al., Intercellular Bioimaging and Biodistribution of Gold Nanoparticle-Loaded Macrophages for Targeted Drug Delivery. Electronics, 2020. 9(7).

51. Nova, Z., H. Skovierova, and A. Calkovska, Alveolar-Capillary Membrane-Related Pulmonary Cells as a Target in Endotoxin-Induced Acute Lung Injury. International Journal of Molecular Sciences, 2019. 20(4).

52. Nasyrova, R.F., et al., Role of nitric oxide and related molecules in schizophrenia pathogenesis: biochemical, genetic and clinical aspects. Front Physiol, 2015. 6: p. 139.

53. van der Vliet, A., J.P. Eiserich, and C.E. Cross, Nitric oxide: a pro-inflammatory mediator in lung disease? Respir Res, 2000. 1(2): p. 67–72.

54. Coleman, J.W., Nitric oxide in immunity and inflammation. Int Immunopharmacol, 2001. 1(8): p. 1397–406.

55. Iverson, N.M., E.M. Hofferber, and J.A. Stapleton, Nitric Oxide Sensors for Biological Applications. Chemosensors, 2018. 6(1).

56. Leslie, K.G., et al., Expanding the Breadth of 4-Amino-1,8-naphthalimide Photophysical Properties through Substitution of the Naphthalimide Core. Chemistry-a European Journal, 2018. 24(21): p. 5569–5573.

57. Y ang, K., et al., Tailoring the properties of a hypoxia-responsive 1,8-naphthalimide for imaging applications. Org Biomol Chem, 2018. 16(4): p. 619–624.

58. Chokshi, N.K., et al., The role of nitric oxide in intestinal epithelial injury and restitution in neonatal necrotizing enterocolitis. Semin Perinatol, 2008. 32(2): p. 92–9.

59. Tian, Y., et al., Quality and efficiency assessment of six extracellular vesicle isolation methods by nano-flow cytometry. J Extracell Vesicles, 2020. 9(1): p. 1697028.

60. Trenchevska, O., R.W. Nelson, and D. Nedelkov, Mass Spectrometric Immunoassays in Characterization of Clinically Significant Proteoforms. Proteomes, 2016. 4(1).

61. Vogel, R., et al., Quantitative sizing of nano/microparticles with a tunable elastomeric pore sensor. Anal Chem, 2011. 83(9): p. 3499–506.

62. Khanal, D., et al., Biospectroscopy of Nanodiamond-Induced Alterations in Conformation of Intra- and Extracellular Proteins: A Nanoscale IR Study. Analytical Chemistry, 2016. 88(15): p. 7530–7538.

63. Cohen, J.M., J.G. Teeguarden, and P. Demokritou, An integrated approach for the in vitro dosimetry of engineered nanomaterials. Part Fibre Toxicol, 2014. 11: p. 20.

64. Xu, F. and F.C. Zhou, Inhibition of microRNA-92a ameliorates lipopolysaccharide-induced endothelial barrier dysfunction by targeting ITGA5 through the PI3K/Akt signaling pathway in human pulmonary microvascular endothelial cells. International Immunopharmacology, 2020. 78.

