## Supplementary data including graphical abstract for "New multiscale characterisation methodology for effective determination of isolation-structure-function relationship of extracellular vesicles"

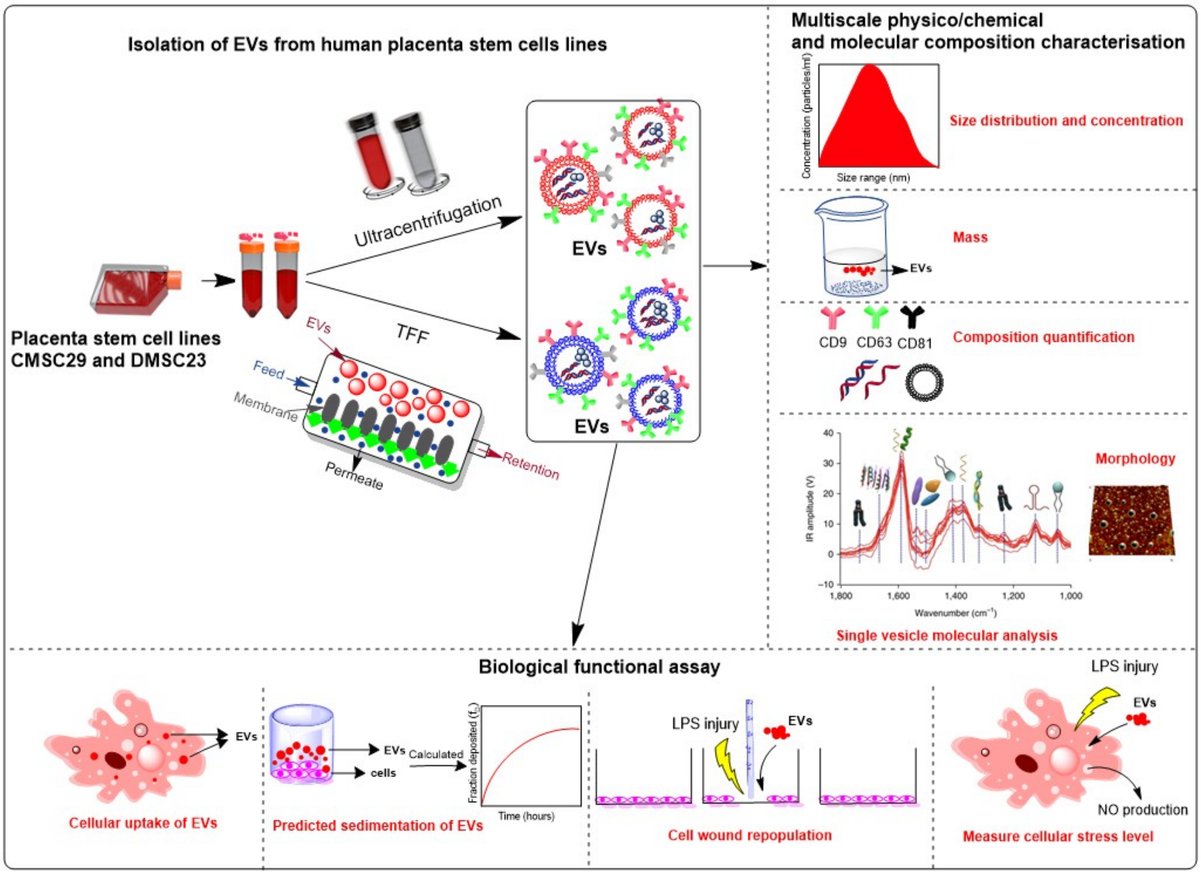


### Supplementary data

#### Isolation of EVs by Exodisc

BT-474 cells were obtained from ATCC (Manassas, VA, USA). EV production medium was prepared by adding exosome-depleted fetal bovine serum (FBS; Thermofisher Scientific, Waltham, MA, USA) to the cell culture medium described in the ATCC protocol. Cells were seeded in T75 cell culture flasks (Corning Inc., Corning, NY, USA) and incubated for 1 day, washed twice in PBS, and 10 mL EV production medium was added to obtain EVs. After 48 h of incubation, EV-containing media was collected and centrifuged at 800 x g for 10 min to remove cells and debris. The supernatant was then injected on an Exodisc (LabSpinner, South Korea) and centrifuged at 3000 rpm (approximately 500 x g) using a bench-top operating machine to separate EVs. EVs separation by Exodisc was done by TFF operated in a centrifugal microfluidic device as described in previous work [64]. The size distribution measured by DLS is given in **Supplementary Fig. S1** (The mean size of EVs was 208.6 ± 69.06 nm and the polydispersity index (PDI) was 0.112)

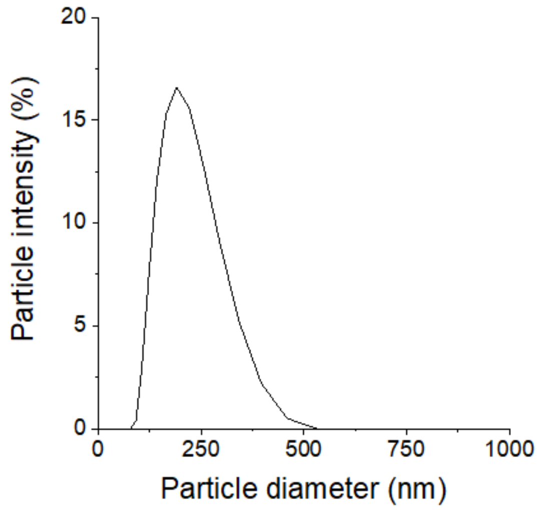


**Fig. S1. The size distribution of the EVs prepared by Exodisc.** The mean size of EVs was 208.6 ± 69.06 nm and the polydispersity index (PDI) was 0.112.

**Table S1.** Summary of averaged particle size and concentration measured using PTA, DLS and nFCM methods for CEVs and DEVs isolated using TFF and ultracentrifugation.

| EV types | **PTA** Mode (nm) | Concentration (EVs/mL) | **DLS** Mode (nm) | **nFCM** Mode (nm) | Concentration (EVs/mL) | **TRPS** Mode (nm) | Concentration (EVs/mL) |
|---|---|---|---|---|---|---|---|
| CEVs TFF | 188 ± 11.5 | $6.1 \times 10^{10}$ | 189 ± 16 | 86.3 ± 19.6 | $9.3 \times 10^{10}$ | 101 ± 3 | $6.71 \times 10^{12}$ |
| CEVs ultra | 161.9 ± 4.7 | $4.63 \times 10^{10}$ | 274 ± 16.7 | 92.9 ± 22.1 | $1.56 \times 10^{11}$ | 90 ± 4 | $1.34 \times 10^{12}$ |
| DEVs TFF | 127.0 ± 4.6 | $3.9 \times 10^{10}$ | 239 ± 24.1 | 85.2 ± 17.1 | $1.92 \times 10^{11}$ | 90.5 ± 6.5 | $5.83 \times 10^{9}$ |
| DEVs ultra | 158.2 ± 9.1 | $4.05 \times 10^{8}$ | 1120 ± 237 | 93.7 ± 21.1 | $2.53 \times 10^{9}$ | 99.5 ± 9.5 | $2.63 \times 10^{8}$ |

**Asymmetric flow-field flow fractionation (AF4)**

To verify the DLS observation of the smaller particles, preliminary AF4 separation and size measurement was carried out. The system used was an Agilent HPLC system (LC1200) (Agilent Technologies Australia, Mulgrave, VIC) coupled to a Wyatt Dawn Helios AF4 system (Wyatt, Santa Barbara, USA). EV suspensions were diluted with ultrapure water or phosphate buffered saline (PBS) and injected into the eluent (water or PBS, respectively), then entered into a flat channel with a 10 kDa membrane and a 350 µm spacer and subjected to a cross-flow gradient (decreasing from 3 mL/min to 0.25 mL/min over 8 min) to facilitate separation. The detector flow was 1 mL/min. The hydrodynamic diameter of the separated EVs were then measured with a multi-angle light scattering system, equipped with a DLS detector positioned at 141 ° to the flow channel. To confirm the hypothesis that the smaller particles were BSA, a sample of pure BSA (Sigma-Aldrich, Australia) was diluted in 10 mM PBS and measured using the same method as the EVs. As shown in **Supplementary Fig. S2**, the elution peak at around 18 minutes coincides for the pure BSA as well as for the CEVs isolated using TFF sample.

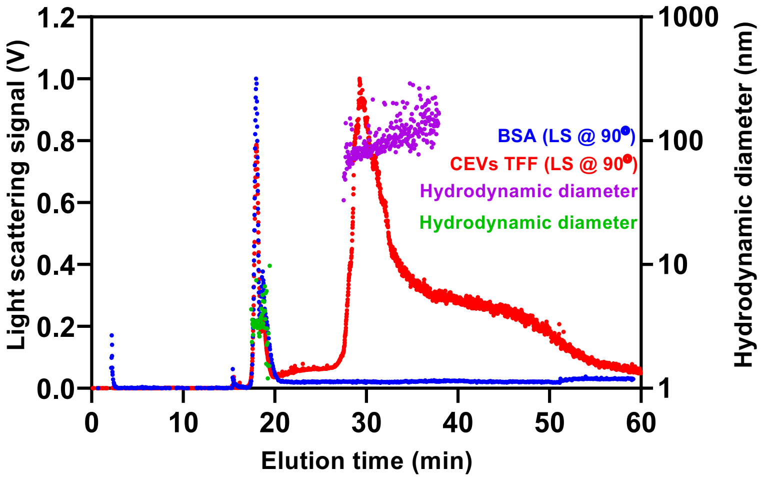


**Fig. S2. A representative AF4 fractionation profile of CEVs isolated using TFF and BSA in PBS** collected by applying a cross-flow gradient with an initial flow rate at 3 mL/min. Light scattering (LS) signal at 90 ° versus elution time.

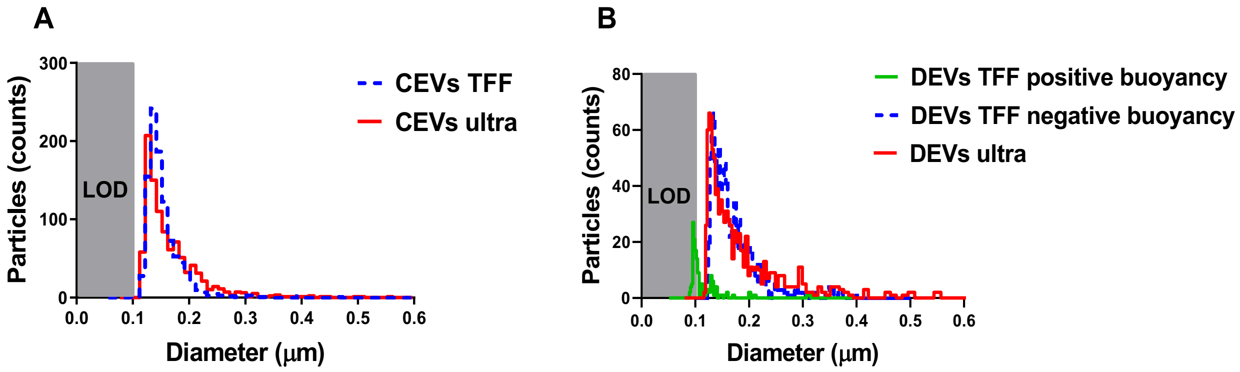


**Fig. S3. Histograms of size distribution of CEVs and DEVs isolated using TFF and ultracentrifugation**. The size distribution based on dry mass of **(A)** CEVs isolated using TFF and ultracentrifugation and **(B)** DEVs isolated using TFF heavier than water, lighter than water and DEVs isolated using ultracentrifugation.

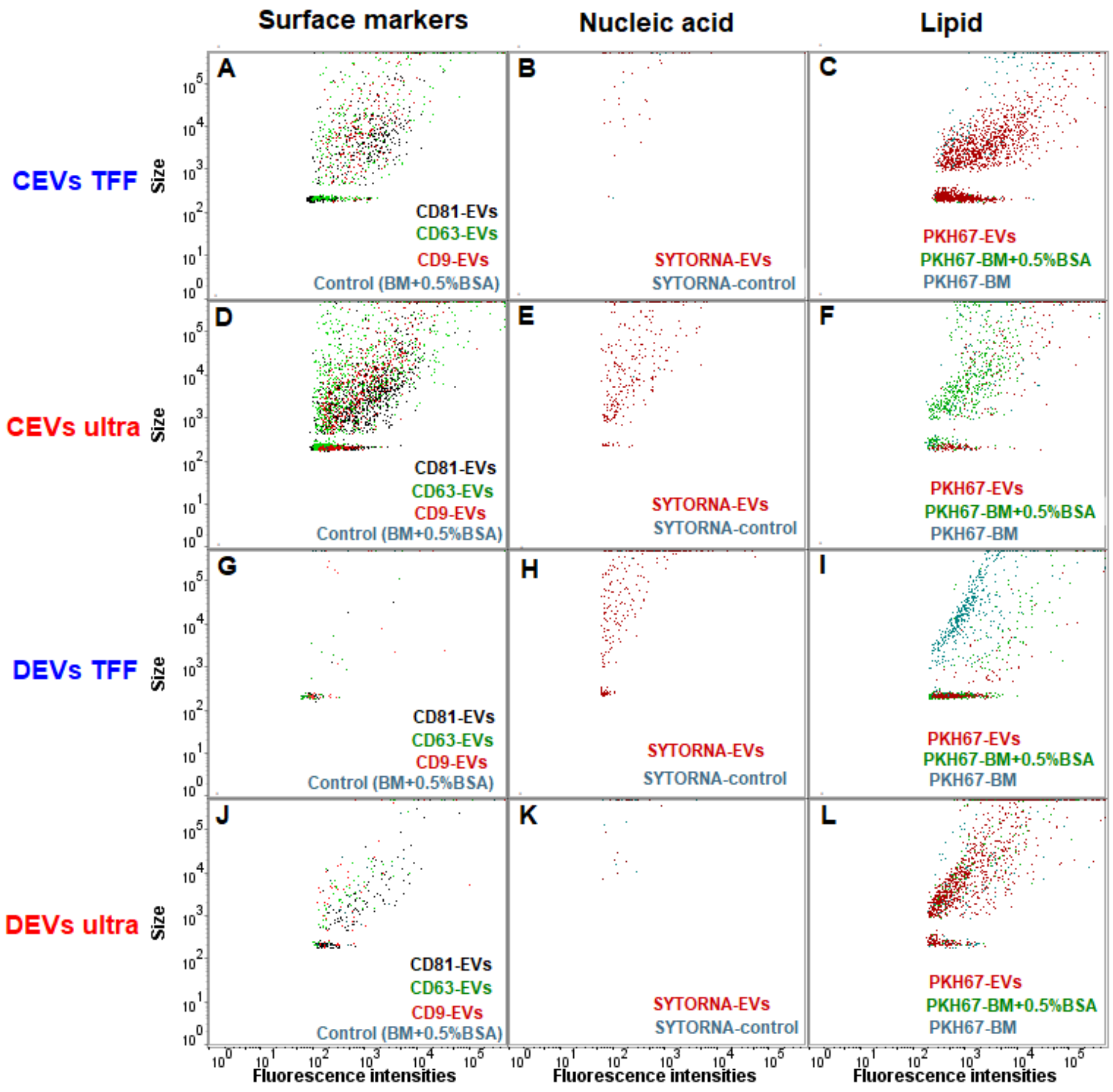


**Fig. S4. The expression level of different surface markers (CD9, CD63, CD81), nucleic acid and lipid of EVs was analysed using nFCM**. Representative dot-plot histograms of CD9, CD63 and CD81 in CEVs isolated using **(A)** TFF and **(D)** ultracentrifugation; DEVs isolated using **(G)** TFF and **(J)** ultracentrifugation. Representative dot-plot histograms of nucleic acid content in CEVs isolated using **(B)** TFF and **(E)** ultracentrifugation; DEVs isolated using **(H)** TFF and **(K)** ultracentrifugation. Representative dot-plot histograms of lipid contents in CEVs isolated using **(C)** TFF and **(F)** ultracentrifugation; DEVs isolated using **(I)** TFF and **(L)** ultracentrifugation.

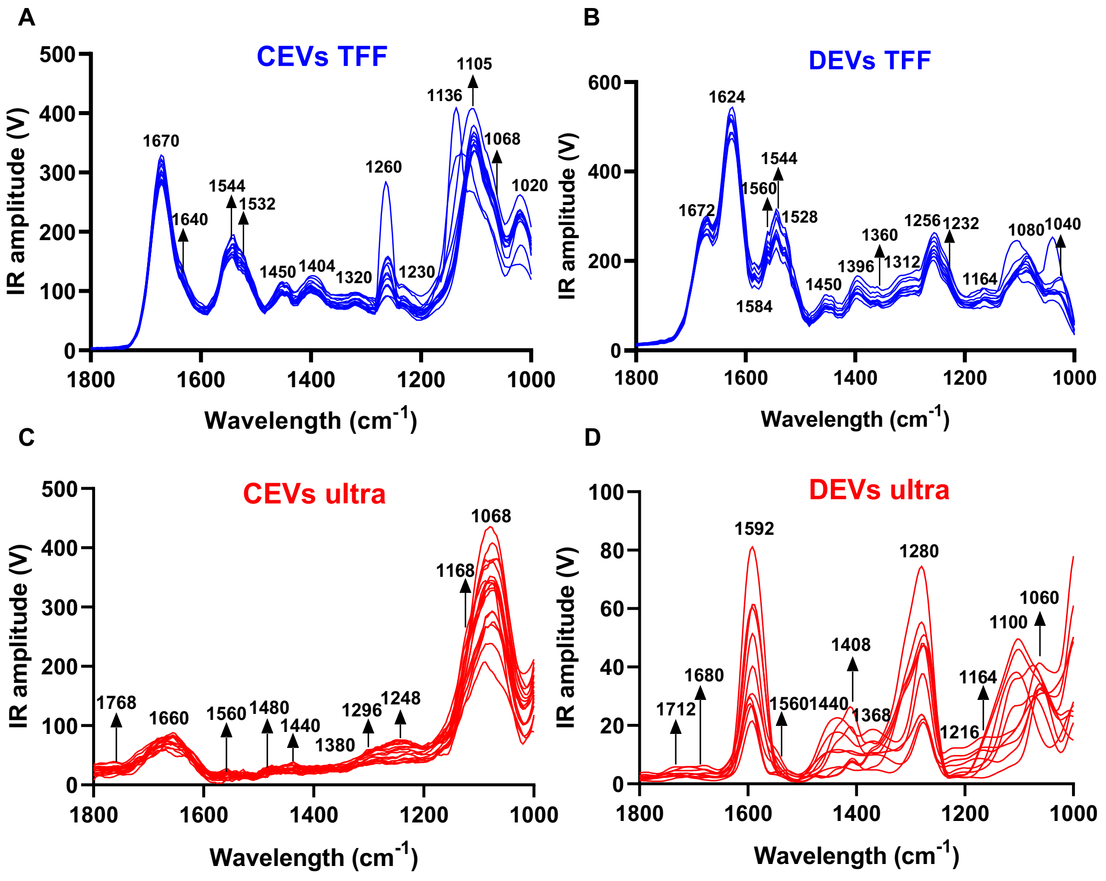


**Fig. S5. Infrared absorption spectra of individual EVs deposited on prism**. AFM-IR spectra of CEVs isolated using **(A)** TFF and **(C)** ultracentrifugation; DEVs isolated using **(B)** TFF and **(D)** ultracentrifugation.

**Table S2.** Parameters used for calculation effective density ($\rho_{effective\ density}$) of CEVs and DEVs isolated using TFF and ultracentrifugation.

| EV samples | $\rho_{media}$ (DMEM + 10% FBS) | $M_{EV}$ | $M_{EVsol}$ (insoluble material) | $V_{pellet}$ (measured in PCV tubes) | SF (stacking factor) | $\rho_{EV}$ |
|---|---|---|---|---|---|---|
| CEVs TFF | 1.0084 | 0.000295 | 0 | 0.00055 | 0.74 | 1.4 |
| CEVs ultra | 1.0084 | 0.000728 | 0 | 0.000225 | 0.74 | 1.4 |
| DEVs TFF | 1.0084 | 0.000749 | 0 | 0.000375 | 0.74 | 1.4 |
| DEVs ultra | 1.0084 | 0.000018 | 0 | 0.0002 | 0.74 | 1.4 |

### Proof of concept for NpNO1 probe

#### Experimental methods

Diethylene glycol monomethyl ether ethylamine was prepared according to a reported procedure [65].

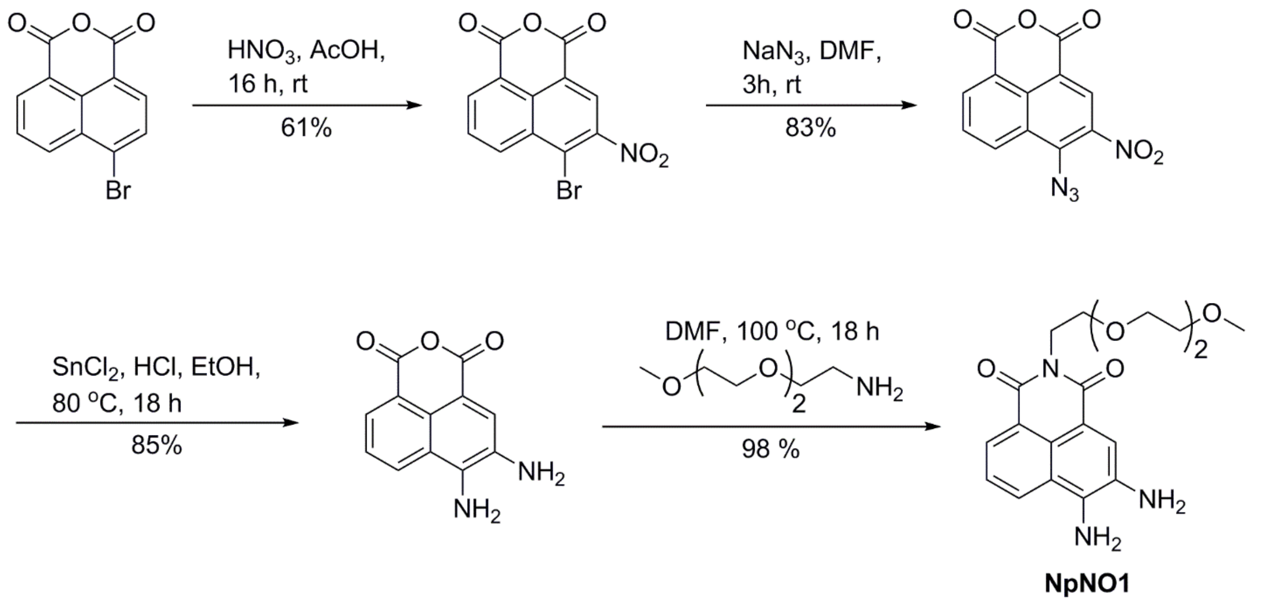


**Scheme S1**: Synthesis of NpNO1

**3-Nitro-4-bromo-1,8-naphthalic anhydride** 4-Bromo-1,8-naphthalic anhydride (5 g, 18.1 mmol) was dissolved in sulfuric acid (35 mL) at 0 °C and sodium nitrate (2 g, 23.52 mmol) added. The reaction mixture was allowed to warm to room temperature and stirred for 16 h, then poured into ice-water (300 mL), and the precipitate filtered to give light yellow solid. The precipitate was recrystallized in glacial acetic acid to afford the product (3.53 g, 61%). **$R_F$** ($CH_2Cl_2$) 0.7; **M.P.** 220-222 °C; **$^1$H NMR** (300 MHz, DMSO-$d_6$): δ 8.92 (s, 1H), 8.84 (d, J = 8.3, 1H), 8.74 (d, J = 6.66, 1H), 8.19 (t, J = 7.8, 1H).

**3-Nitro-4-azido-1,8-naphthalic anhydride** $NaN_3$ (912 mg, 14 mmol) was added to a solution of 3-nitro-4-bromo-1,8-naphthalic anhydride (3 g, 9.31 mmol) in DMF (35 mL) stirred for 3 h at room temperature. The solution was poured into 250 mL of ice-water, and the precipitate was filtered, washed with cold water and cold acetic acid, then dried to give the product as a light brown solid (2.212 g, 83%). **$R_F$** ($CH_2Cl_2$) 0.6; **M.P.** 233-235 °C; **$^1$H NMR** (300 MHz, DMSO-$d_6$): δ 8.89 (m, 2H), 8.70 (d, J = 6.39, 1H), 8.06 (t, J = 7.88, 1H).

**3,4-Diamino-1,8-naphthalic anhydride** 3-Nitro-4-azido-1,8-naphthalic anhydride (0.5 g, 1.76 mmol) was added to a mixture of $SnCl_2$ (3.38 g, 15 mmol) in concentrated hydrochloric acid (10 mL) and stirred for 40 minutes at 50 °C. Ethanol (10 mL) was then added and the mixture stirred at 80 °C for another 18 h. The reaction mixture was then cooled to room temperature, and the resulting precipitate filtered, washed with water, and dried to give the desired product as a red solid (340 mg, 85%). **$R_F$** (DCM 9:1 MeOH) 0.64; **M.P.** >260 °C; **$^1$H NMR** (300 MHz, DMSO-$d_6$): δ 8.59 (d, J = 8.03, 1H), 8.22 (d, J = 6.56, 1H), 7.9 (s, 1H), 7.63-7.56 (m, 1H), 6.87 (s, 2H); **ESI-MS** $(M - H)^-$ 227, $(M + Na)^+$ 251, $(2M + Na)^+$ 479.

**Mesyl-triethylene glycol monomethyl ether** Triethylamine (6.5 mL, 47 mmol) was added to a solution of triethylene glycol monomethyl ether (4.9 mL, 30.5 mmol) in dry dichloromethane (25 mL) under a $N_2$ atmosphere at 0 °C. Mesyl chloride (2.85 mL, 36.5 mmol) in dry dichloromethane (25 mL) was added to the mixture dropwise over 90 min. The mixture was stirred at 0 °C under for

1 h, then at rt for 15 h. The reaction mixture was washed with aqueous HCl (3%) (50 mL) and brine (50 mL) then dried with sodium sulfate and evaporated under reduced pressure to give mesyl-triethylene glycol monomethyl ether as a yellow oil (5.59 g, 76%.) **R$_F$** (PE 5:5 EtOAc) 0.29; **$^1$H NMR** (200 MHz, $CDCl_3$): δ 4.31 (m, 2H / -SO-CH2-), 3.7 (m, 2H), 3.58 (m, 6H), 3.48 (m, 2H), 3.3 (s, 3H), 3.0 (s, 3H).

**N-TEG-3,4-amino-1,8-naphthalimide (NpNO1)** A solution of 3,4-diamino-1,8-naphthalic anhydride (0.1 g, 0.44 mmol), mesyl-triethylene glycol monomethyl ether (80 mg, 0.48 mmol) and DIPEA (100 µL, 0.57 mmol) in DMF (10 mL) was stirred at reflux for 18 h. The DMF was evaporated at 45 °C under reduced pressure and crude residue then purified by silica flash chromatography (5-10% MeOH in $CH_2Cl_2$) to give **NpNO1** as a purple solid (160 mg, 98%). **R$_F$** ($CH_2Cl_2$ 9:1 MeOH) 0.2. **$^1$H NMR** (400 MHz, DMSO-d$_6$): δ 8.49 (dd, $J$ = 8.4, 0.8, 1H), 8.18 (dd, $J$ = 7.2, 0.8, 1H), 7.92 (s, 1H), 7.54 (dd, $J$ = 8.4, 7.2, 1H), 6.51 (s, 2H), 5.16 (s, 2H), 4.19 (t, $J$ = 6.5, 2H), 3.60 (t, $J$ = 6.6, 2H), 3.55-3.42 (m, 8H), 3.17 (s, 3H); **$^{13}$C NMR** (125 MHz, DMSO-d$_6$): δ 164.5, 163.6, 136.9, 131.0, 128.5, 127.7, 124.1, 123.8, 122.0, 121.3, 120.1, 108.7, 71.7, 70.1, 70.1, 70.0, 67.6, 58.4, 38.7. **ESI-MS** $(M + Na)^+$ 396.

**Fluorescence spectroscopy**

All fluorescence spectra were recorded on a Varian Cary Eclipse fluorometer in 1 cm pathlength quartz cuvettes.

**Cell culture and microscopy**

A549 (adenocarcinoma human alveolar basal epithelial) cell line was maintained in exponential growth as monolayers in Advanced Dulbecco's Modified Eagle's Medium (ADMEM) supplemented with 2 mM glutamine and 2 % foetal bovine serum (FBS). Cells were incubated at 37.0 °C in 5 % (v/v) $CO_2$ under humidified conditions. The cell line was passaged using 0.25 % trypsin to facilitate dislodgement of cells from the flask.

Approximately 1 x $10^4$ DLD-1 cells were seeded on a poly-D-lysine coated MatTek® dish and allowed to adhere overnight. The adhered cells were stained with 50 µM NpNO1 with or without 5 mM MAHMA NONOate (the NO donor) in ADMEM (supplemented with 2 mM glutamine and 2 % FBS) for 24 h at 37 °C, 5 % (v/v) $CO_2$ under humidified conditions. The cells were then washed three times with phosphate-buffered saline (PBS) and imaged in FluoroBrite™ media (supplemented with 2 mM glutamine and 2 % FBS).

Z-stacked confocal micrographs with a 1.0 µm step-size and a resolution of 0.207 µm/pixel were acquired using the Olympus FluoView® FV3000 confocal microscope. The microscope is equipped with a UPLSAPO 60X water-immersion objective lens (numerical aperture = 1.15), connected to a computer installed with the FV10-ASW viewer software v1.7 (Olympus). Cells were visualised using the 405 nm laser as an excitation source, and emission between 415-515 nm were acquired by the detectors (PMTs) of the microscope. Acquired micrographs were processed and analysed using ImageJ FIJI (1.49u, Java 1.6.0_24).

**Statistical analyses**

Statistical tests were carried out using GraphPad Prism® software (version 7.02; GraphPad, LaJolla, CA, USA). Multi-group comparisons were made using one-way analysis of variance (ANOVA) with an alpha of 0.05 [66]. A $P$-value of $<0.05$ was regarded as statistically significant. If the null hypothesis is rejected ($P < 0.05$), a *post-hoc* Tukey test was performed for pairwise comparisons. Results presented were as the mean values ± standard error of the mean, unless otherwise stated.

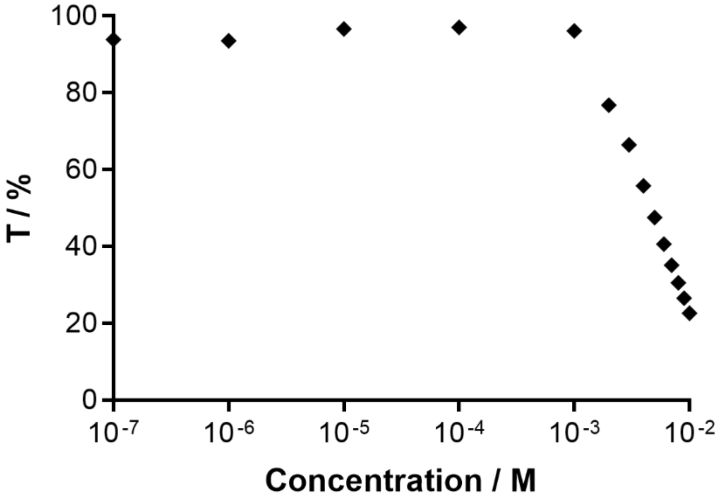


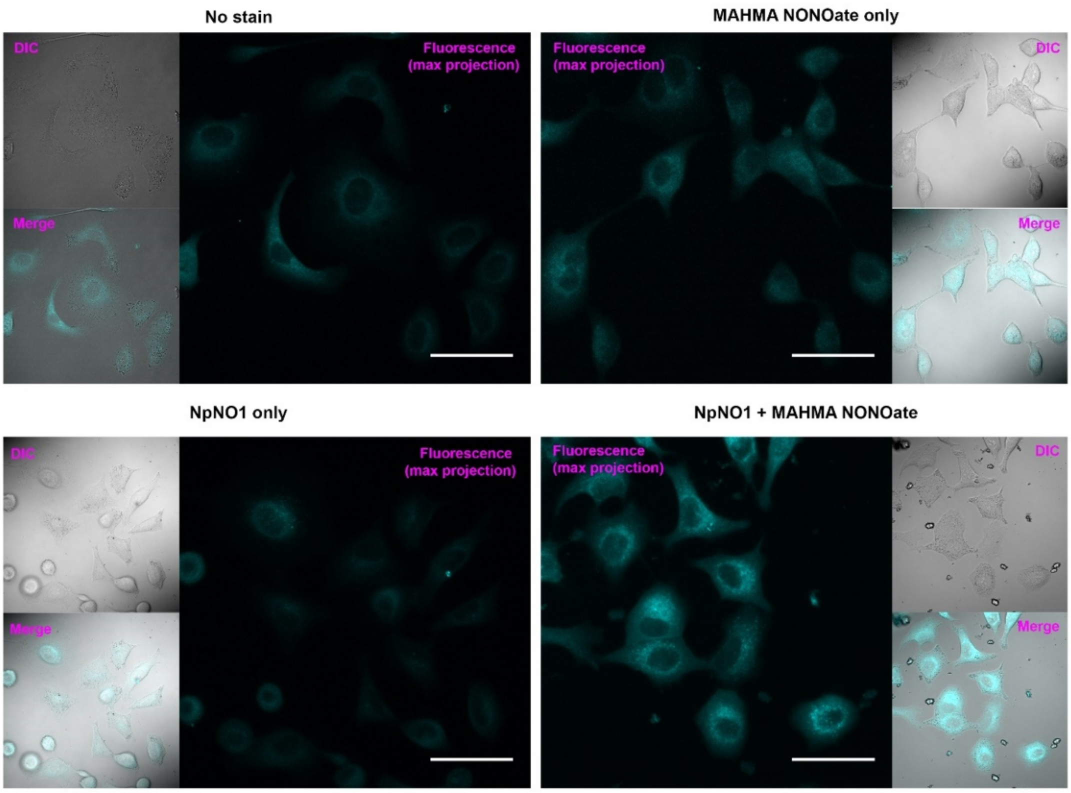


**Fig. S6. Normalised transmittance at 650 nm of NpNO1 at increasing concentrations and representative micrographs of A549 cells treated in various conditions**. Z-stacked confocal micrographs were acquired, and maximum projection of the z-stacks are shown. The fluorescence micrographs were accompanied with respective differential interference contrast (DIC) images and merged images. Scale bar represents 50 μm.

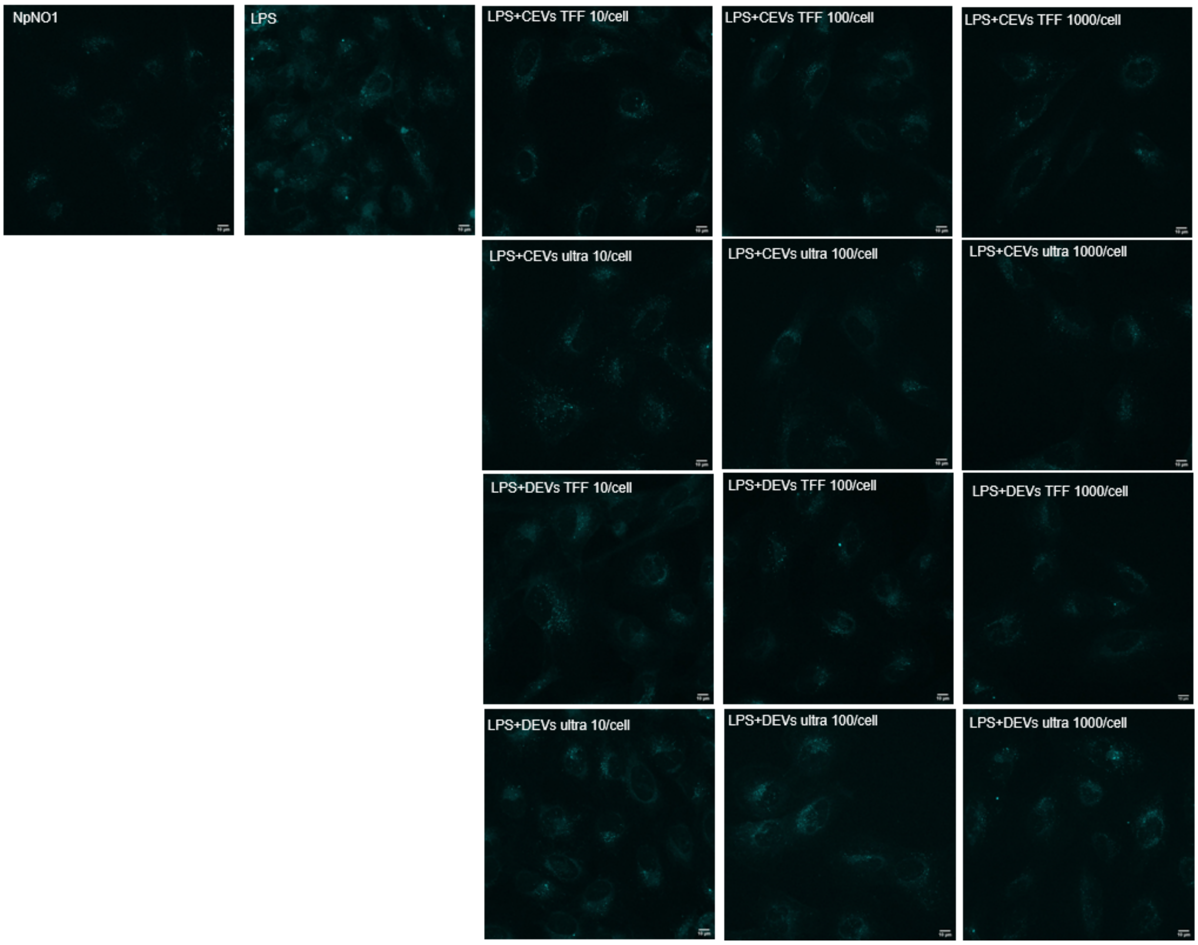


**Fig. S7. Representative micrographs of BEAS-2B cells treated with various EV concentrations.** Maximum projection of the z-stacks is shown. Scale bar represents 10 µm for each condition.
